# Whole genome sequencing-based multi-locus association mapping for kernel iron, zinc and protein content in groundnut

**DOI:** 10.1101/2025.08.04.668427

**Authors:** U. Nikhil Sagar, Sejal Parmar, Sunil S. Gangurde, Vinay Sharma, Arun K. Pandey, D. Khaja Mohinuddin, Namita Dube, Ramesh S. Bhat, K. John, Muga D. Sreevalli, P. Sandhya Rani, Kuldeep Singh, Rajeev K. Varshney, Manish K. Pandey

**Affiliations:** Center of Excellence in Genomics & Systems Biology (CEGSB) and Center for Pre-Breeding Research (CPBR), International Crops Research Institute for the Semi-Arid Tropics (ICRISAT), Hyderabad-502324, Telangana, India; Department of Genetics and Plant Breeding, S.V. Agricultural College, ANGRAU, Tirupati-517502, India; Discipline of Biotechnology, University of Agricultural Sciences, Dharwad-580005, India; Centre for Crop and Food Innovation, WA State Agricultural Biotechnology Centre, Murdoch University, Murdoch, Australia

**Keywords:** Candidate genes, Fe and Zn homeostasis pathway, KASP markers, Marker trait associations, Nutritional quality

## Abstract

Malnutrition is a major global challenge, especially in the developing regions, where improving the nutritional content of staple crops is a significant step towards alleviating hidden hunger. Groundnut, a nutrient rich legume contains several mineral nutrients, high protein, essential amino-acids and vitamins that are required for human health. In this study, multi-season phenotyping data for kernel iron (Fe), zinc (Zn) content and protein content (PC) and whole genome re-sequencing (WGRS) data on mini-core collection, were used to perform genome-wide association study (GWAS) analysis. Phenotypic variability analysis revealed a large variation in the Fe (7.6 – 42.8 ppm), Zn (10.9 – 62.4 ppm) and PC (12.7 – 33.6%). GWAS analysis identified a total of 15 marker-trait associations (MTAs) and 28 candidate genes for pooled season data, as well as 44 MTAs and 62 candidate genes for individual seasons. Key candidate genes like *MYB transcription factor* (*Arahy.QI0PHV*, *Arahy.1I6ZSS*), *Zn finger MYM type protein*, *RING finger MYM type protein* (*Arahy.7P97F6*, *Arahy.9R964H*, *Arahy.I3B88T*) and *NAC domain protein* (*Arahy.LV3APC*), associated with the Fe and Zn homeostasis pathway, genes related to protein homeostasis, such as *protein kinase family protein* (*Arahy.4D7KBI*), and *E3 ubiquitin-protein ligase* (*Arahy.PE3CF6*), were also identified within significant MTA regions. These findings provide basis for the detection and characterization of the possible candidate genes related to the nutritional quality traits. Single nucleotide polymorphism (SNP)-based KASP (Kompetitive Allele Specific Polymerase Chain Reaction) markers for 9 SNPs were designed and validated. Of these, three markers (snpAH00636, snpAH00641 and snpAH00644) showed polymorphism which could be deployed in the genomics-assisted breeding for the development of nutrient-rich groundnut varieties.

## 1. INTRODUCTION

Groundnut (*Arachis hypogaea* L.), is often referred to as the “King of oilseeds” due to its high oil content, nutritional value, and economic importance. It is also referred to as peanut, earthnut, monkey nut, wonder nut, and goobernut (1). It is the fourth-most significant vegetable protein source in the world. This self-pollinated, segmental allotetraploid crop with minimal molecular-level variation (2) belongs to the family *Leguminosae* and sub-family *Papilionaceae* (3). While it originated in South America-it is now cultivated worldwide across tropical, sub-tropical, and semi-arid regions. Usually it is grown on over 32.7 million hectares across more than 100 countries, producing approximately 53.9 million tonnes ha^-1^ with a productivity of 1.64 tonnes ha^-1^ (Food and Agriculture Organization (FAO), 2023) [https://www.fao.org/faostat/en/#data]. China and India lead global production of groundnut, followed by USA, Sudan, and Nigeria.

Fe and Zn are the vital micronutrients needed for human growth and development, as they serve as cofactors for proteins like haemoglobin and cytochromes (4). Fe and Zn deficiencies cause anaemia—also known as “hidden hunger” or malnutrition—affects nearly three billion people globally (World Health Organization, 2016), a considerable proportion of women and children in India suffer from anaemia. Groundnut, as a nutrient rich legume crop, contains 25% to 28% protein, 48% to 50% oil, and 10% to 20% carbohydrates (4). Moreover, it is also a valuable source of monounsaturated and polyunsaturated fatty acids, proteins and carbohydrates. The seeds are rich in bioactive polyphenols, flavonoids, isoflavones, and several B-complex vitamins, and antioxidants, including resveratrol and p-coumaric acid. Raw groundnut contains 4.58 mg of Fe, 3.27 mg of Zn and 26 g of PC per 100g. Groundnuts and their products can be marketed as nutrient-dense foods to address protein, energy, and micronutrient deficits in vulnerable groups because of their high nutritional content (6). For instance, ready-to-use therapeutic products (RUFTs) made from groundnuts, such as plumpy nuts, have already proven to be successful in saving the lives of malnourished children in certain regions (7).

The likelihood of advanced genomic resources in groundnut-such as genotyping assays, reference genomes, and diagnostic markers (8) has opened up new opportunities for their use in genomics and breeding to enhance its nutritional profile. High-to-mid-density genotyping arrays/assays were used for constructing genetic maps, identifying genomic regions and candidate genes via GWAS, and designing diagnostic markers to accelerate breeding programs (4). Further, these assays provide potentiality for identifying varietal seed mixtures, assessing genetic purity in seed systems and gene bank germplasms, conducting foreground and background selection in backcross breeding, facilitating trait mapping, and genomic selection in groundnut (9). GWAS is one of the most powerful methods as it helps in improving statistical power or resolving computation complexity problems in a wide range of mapping populations including natural populations with broad genetic base and breeding populations with narrow genetic base. It also helps in the detection of candidate genes linked with complex traits, and also provides higher accuracy in mapping and detecting associations between molecular markers and key traits (10).

Several key candidate genes carrying transcription factors such as *NAC*, *MYB* were found to be regulating seed and pod development, plant growth, and photosynthesis, under the Fe deficiency conditions (11). The proteins encoding *MYB transcription factor*, *RING finger protein*, and *NAC transcription factor* were reported to have a key role in Fe and Zn homeostasis pathways in the seeds (4). Earlier studies reported numerous potential homologs responsible for the transport of Fe from leaf to the root, such as *IRT1*, *FIT1*, *bZIP23*, and *OPT3*, identified in groundnut and other legume crops (12). Previous studies in cereals identified several genes or gene families that are responsible for Fe and Zn homeostasis including *YSL*, *NRAMP*, *ZIP*, *Heavy metal ATPase*, *cation diffusion facilitator family*, *nicotianamine synthase*, and *nicotianamine aminotransferase* (13). The primary objective of this study was to identify the MTAs associated with Fe, Zn and PC, and to pinpoint the candidate genes required for the homeostasis of Fe, Zn and PC in groundnut seeds. In this context, the WGRS genotyping data using 5,66,320 polymorphic SNPs for the mini-core set, combined with phenotypic data generated for Fe, Zn and PC was used in the GWAS. The candidate genes encoding various important proteins are known to be regulating the homeostasis of Fe, Zn and PC and are used for the development and validation of molecular markers.

## 2. MATERIALS AND METHODS

### 2.1. Multi-environment and multi-location phenotyping for Fe, Zn and PC

The groundnut mini-core set comprising 184 lines (14) was used to conduct the experiment. The experiment was carried out at ICRISAT, Patancheru, India for five seasons; post-rainy 2009-10 (S1), rainy 2009 (S2), post rainy 2010-11 (S3) as reported in Upadhyaya et al., (2012) (15), rainy 2020 (S4) and rainy 2021 (S5) for estimation of Fe and Zn contents (ppm) (**Supplementary Table S1**). The PC data was estimated from two geographic locations, ICRISAT during the four seasons-rainy 2008, rainy 2009 as reported in Upadhyaya et al., (2012) (15), rainy 2020 and rainy 2021 and Dharwad during the two rainy seasons of 2008 and 2009 as reported in Pandey et al., (2014) (16). The frequency distribution graphs (violin plots), Pearson correlation plot and Principal component analysis (PCA) of Fe, Zn and PC were generated using “SRplot” (17).

Fe and Zn contents were determined using the ICP-OES method (18). The phenotypic data for Fe and Zn contents during S1, S2, and S3 were estimated at Charles Renard analytical laboratory, ICRISAT, whereas the Fe and Zn contents during S4 and S5, were estimated at NCML laboratory, Hyderabad. Approximately 0.3 g of oven-dried samples were prepared by digesting these overnight in 2 ml nitric acid and 0.5 ml hydrogen peroxide, followed by a series of heating steps and final dilution to 25 ml. The filtered supernatant was analyzed using ICP-OES (4).

The PC (%) was quantified using Near infrared-reflectance spectroscopy (NIRS). Samples from the two rainy seasons of 2008 and 2009, collected at ICRISAT and Dharwad, were analyzed using NIRS. The estimation of PC during the two rainy seasons, 2020 and 2021 was done at NCML laboratory, Hyderabad. The groundnut samples (approximately 60 g) were scanned with a diode array analyzer at wavelengths from 950 to 1650 nm. Absorbance readings were recorded in 5 nm increments, and each sample was scanned three times to ensure accuracy (4).

### 2.2. Genomic DNA extraction, sequencing and SNP calling

The nucleospin plant II kit was used to extract the genomic DNA from 100 mg of leaf sample tissue. 500 μL of lysis buffer was used to homogenize the leaf sample, and then 10 μL RNAse was added to remove the RNA impurities. The samples were incubated at 65°C for 1 hour in water bath, then centrifuged at 5500 rpm for 10 min. A separate tube was used to collect the supernatant, and binding buffer of 450 μL was added. This whole mixture was centrifuged at 6000 rpm for 1 min after being filtered using a NucleoSpin plant MN column. In the final step, pre-heated elution buffer of 50 μL was added to the column’s membrane filter, incubated at 65°C for 5 minutes, and centrifuged at 6000 rpm for 1 min to elute the DNA (19). 0.8% agarose gel was used to check the DNA quality whereas the quantity of DNA was assessed using NanoDrop 8000 Spectrophotometer.

The TruSeq library kit was used to generate paired-end sequencing libraries with a 500 bp insert size to represent each genotype. Short reads (150 bp) with paired ends were generated by the Illumina HiSeq 2500 platform. Sequence data is provided with bio-project ID (PRJNA1002116, PRJNA490835 and PRJNA490832). The adapter sequences were initially cut after post-sequencing, and then the low-quality reads i.e., those with > 20% low-quality bases (quality value ≤ 7) and > 5% “N” nucleotides—were filtered out using SOAP2 (20) to produce high-quality sequencing data for further analysis. The clean sequence reads were aligned to the reference genome of the cultivated tetraploid “Tifrunner” with the parameters “-m 300 -x 600 -s 35 -l 32 -v 5 -p 4”. SOAPsnp3 was used to estimate the likelihood of each potential genotype for every sample in order to determine the maximum likelihood estimation of the frequency of alleles in the population. Variants of low quality were filtered based on strict filtering criteria, including sequencing depth (> 10,000 and < 400), mapping times (> 1.5), and a quality score (<20). SNP loci were determined with an estimated allele frequency neither equal to 0 nor 1. Furthermore, SNPs with 50% or more missing data among genotypes were also removed, resulting in a set of high-quality SNPs for downstream analysis.

### 2.3. GWAS for Fe, Zn and PC

GWAS analysis was conducted using multi-season phenotyping data from 184 mini-core set and SNPs called using WGRS data. The Genome Association and Prediction Integrated Tool (GAPIT) package (21) in R software was employed to identify significant MTAs. In this study, several popular GWAS models like BLINK, MLMM, SUPER, CMLM, FarmCPU, GLM and MLM were used. The significant MTAs were found using single locus models-the Settlement of MLM Under Progressively Exclusive Relationship (SUPER), the Mixed Linear Model (MLM), the Compressed MLM (CMLM), and the General Linear Model (GLM), and multi-locus models-Multiple Loci Mixed Linear Model (MLMM), Bayesian-information and Linkage-disequilibrium Iteratively Nested Keyway (BLINK) (22) and the Fixed and Random Model Circulating Probability Unification (FarmCPU) (23). The FarmCPU method increases statistical power by controlling false positives, removing confounding, and splitting Multiple Loci Linear Mixed Models into random effects model (REM) and fixed effects model (FEM). The SUPER model handles MLM’s computational problems, and BLINK uses LD information to increase statistical power (24).

Among the various models used for GWAS of pooled data, the threshold level of ‘*p*’ was set to 6. The Manhattan plots depicted the output of GWAS and the MTAs at a significant threshold level of above 6 were considered as significant MTAs. For seasonal analysis, the threshold level of ‘*p*’ was set to 6.7. Additionally, the Q-Q plots were used to select these markers based on their elevated values indicating their non-random association with the trait.

The Q-Q plots were used to determine the best model for each trait. The significant MTAs were identified using the Bonferroni correction, which is defined as the significance level divided by the number of markers at each locus e.g. 0.05/number of markers (25). The Q-Q plots measured by different models represent the comparison between the observed and expected log_10_ (*p-value*) to identify the significant SNP loci linked with the Fe, Zn and PC. The trait-related genomic regions were mined using the Bonferroni-corrected *p-value*, and the markers that had a *p-value* of 0.05/N (N = total number of SNPs) or less were considered significant.

### 2.4. Identification of candidate genes from MTAs for Fe, Zn and PC

Candidate genes linked with the significant MTAs were discovered using peanut base (https://Peanutbase.org/) through GBrowse (cultivated peanut) version 1. Trait-associated genomic regions were examined, and candidate genes with known roles in Fe, Zn, and PC homeostasis pathways were highlighted. These genes serve as a valuable resource for the development of molecular markers and for further genetic and functional characterization (26).

The expression of the candidate genes was examined using the *Arachis hypogaea* gene expression atlas (AhGEA), which was created for the subspecies *hypogaea* (27). Expression data were analysed for 16 tissues (Pre-soaked seeds, emerging radicle, cotyledons, embryo, root seedling, leaves veg, root veg, shoot veg, shoot seedling, immature bud, seeds_05, seeds_15, seeds_25, pod wall immature, pod wall mature, nodules), providing insights into functional relevance of candidate genes in homeostasis of Fe, Zn and PC.

### 2.5. KASP marker development and validation

SNPs influencing the candidate gene functions were selected for the development of KASP markers using sequences from the 300-bp upstream and 300-bp downstream regions (28) (**Supplementary Table S2**). For each SNP marker, KASP assays were designed using two allele-specific forward primers and one common reverse primer. The SNPs associated with Fe, located on 6 different chromosomes-Ah03, Ah05, Ah14, Ah18, Ah19, Ah20 and the SNP associated with Zn, located on chromosome Ah05 were selected for the development of markers. These KASP markers were used for validation in diverse germplasm lines of ICRISAT comprising 30 high Fe and Zn content lines, 20 low Fe and Zn content lines.

## 3. RESULTS

### 3.1. Phenotypic variability, correlation and multi-variate association among Fe, Zn and PC

The phenotypic distributions of Fe, Zn and PC across multiple seasons and locations were visualized using violin plots (**Figure 1A**), that showed a wide variability. The mean value of Fe varied across different seasons as 29.8 ppm for S1, 21.9 ppm for S2, 23 ppm for S3, 26.7 ppm for S4 and 19.6 ppm for S5. Similarly, mean Zn showed a range of 43 ppm in S1, 27.7 ppm in S2, 34.8 ppm in S3, 35.5 ppm in S4 and 27.2 ppm in S5. Likewise, the PC also exhibited considerable variation, with mean values of 25.1% for 2008 rainy, 22.8% for 2009 rainy season at ICRISAT and 23.9% for 2008 rainy, 23.3% for 2009 rainy season at Dharwad, 23.6% for S4 and 22.7% for S5 at ICRISAT.

**Figure 1.**
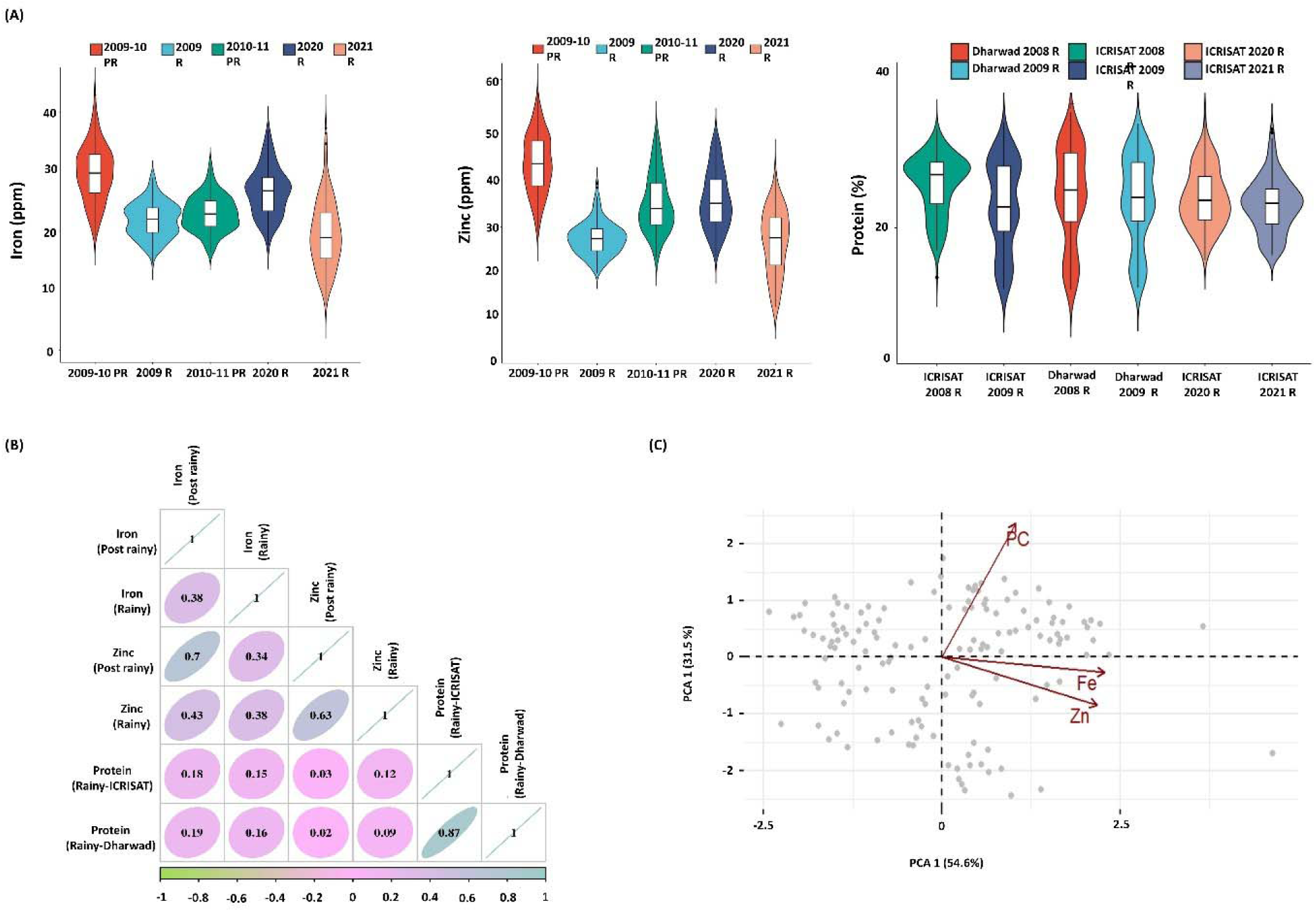
Phenotypic variation for Fe, Zn and PC in the diverse mini-core set. (A) Violin plots showing the distribution of Fe (ppm), Zn (ppm), and PC (%) for various genotypes across different years under both rainy (R) and post-rainy (PR) seasons. The box plot represents the average frequency of distribution of genotypes in a particular season (B) The correlation matrix indicating relationships among Fe, Zn and PC across seasons and locations, with significant positive correlation between Fe and Zn in pooled post-rainy season (*r* = 0.7) (C) Principal component analysis (PCA) demonstrating the distribution of the groundnut accessions (each dot on the plot represents an accession), based on the first two principal components, with PC1 accounting for 54.6% of the variance and PC2 accounting for 31.5%, cumulatively explaining 85.9% of the total variability.

Across the seasons, Zn was consistently higher than the Fe which ranged from 7.56 ppm (ICG 2925) in S5 to a maximum of 58.22 ppm (ICG 5827) in the S4 whereas the Zn showed a minimum range of 13.74 ppm (ICG 4412) in S5 and a maximum range of 62.35 ppm (ICG 13856) in S4. The PC varied from a minimum value of 12.74% (ICG 3775) in 2008 rainy season at Dharwad to a maximum of 33.59% (ICG 2773) in S4 at ICRISAT.

Correlation analysis among Fe, Zn and PC (**Figure 1B**) showed a significant positive correlation between these traits. The significance of correlations was considered at a level of *p-value* = 0.05. The strongest correlation was observed between Fe in pooled post rainy and Zn (*r* = 0.7) in pooled post rainy followed by Fe in pooled post rainy with Zn in pooled rainy (*r* = 0.43). Partial but significant positive associations were also observed between Fe in pooled rainy and PC in pooled rainy at ICRISAT (*r* = 0.15), followed by Fe in pooled rainy and PC in pooled rainy at Dharwad (*r* = 0.16). Similar associations were observed between Zn in pooled rainy and PC in pooled rainy at ICRISAT (*r* = 0.12), and between Zn in pooled rainy and PC in pooled rainy at Dharwad (*r* = 0.09).

For multivariate analysis, PCA is a very useful technique to determine the key factors from complex traits that show significant correlation and also covers eigen vectors, covariance and standard deviation (29). In this study, three traits were used to perform the PCA and each of these traits contributed to some amount of variance in the two principal components. The scatter plot (**Figure 1C**) depicts the distribution of genotypes into two subgroups, revealed variations within these subgroups. The first principal component (PC1) (x-axis) accounted for 54.6% of total variation, while the second principal component (PC2) (y-axis) explained 31.5%, together accounting for 86.1% of the total variance. The Fe and Zn across all the seasons contributed positively towards the PC1, while PC contributed positively to both PC1 and PC2. The Fe and Zn contents were nearly parallel to each other, indicating a strong positive correlation, whereas, the PC is oriented differently from both the Fe and Zn, indicating a weaker positive correlation.

### 3.2. Identification of MTAs

GWAS analysis identified 15 significant MTAs for the three traits (4 MTAs for Fe, 4 for Zn, and 7 for PC) (**Table 1**). The high number of pooled MTAs were observed on chromosome Ah17 (3 MTAs), followed by 2 MTAs each on chromosomes Ah02, Ah13, and Ah19. For seasonal data analysis, a total of 44 significant MTAs were observed, with 20 MTAs associated with Fe, 13 for Zn and 11 for PC (**Table 2**). The maximum number of MTAs were identified on chromosome Ah03 (8 MTAs), followed by Ah17 (5 MTAs) and 4 MTAs each on chromosome Ah05 and Ah19.

**Table 1.**
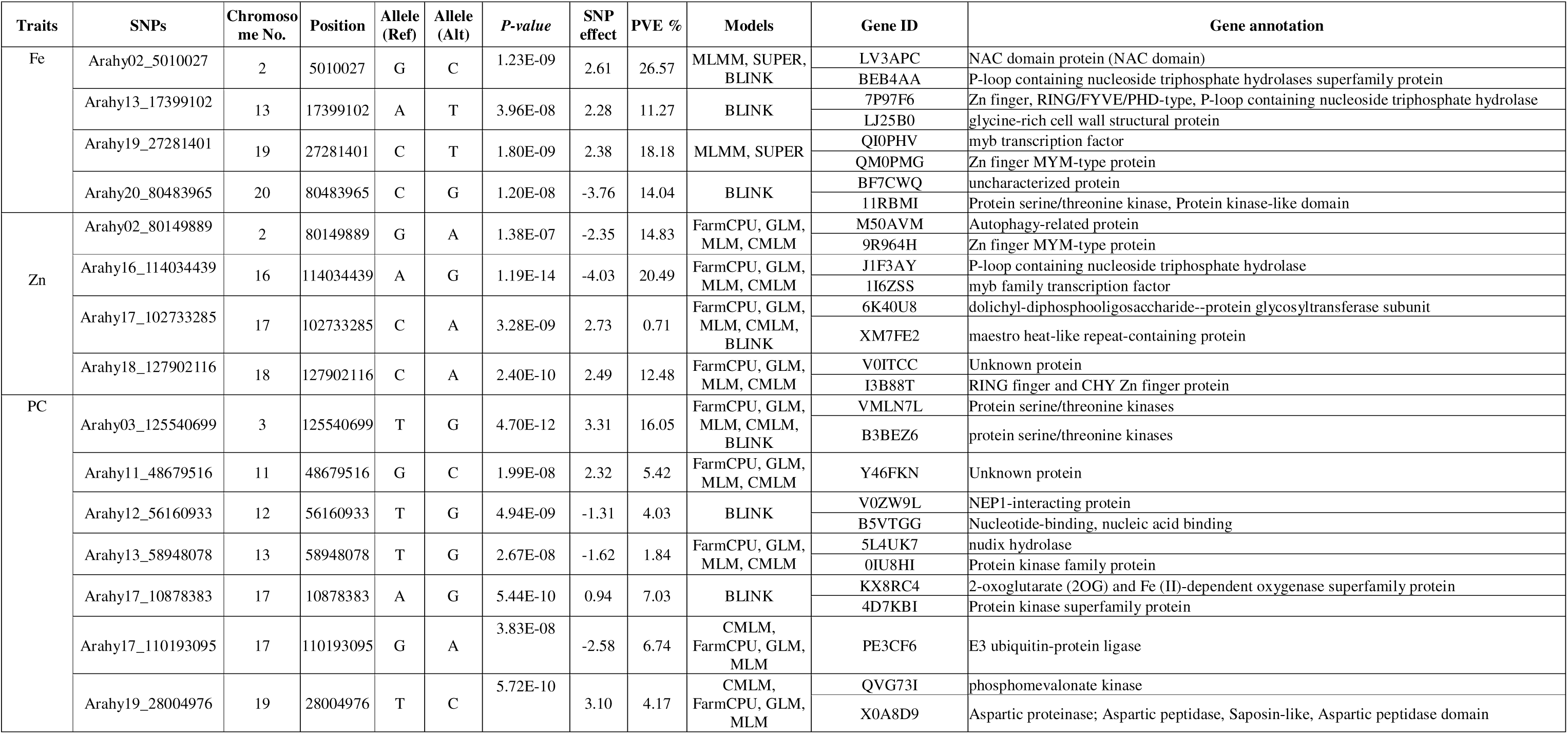
List of candidate genes in significant MTAs for Fe, Zn and PC using different models for pooled season data.

**Table 2.**
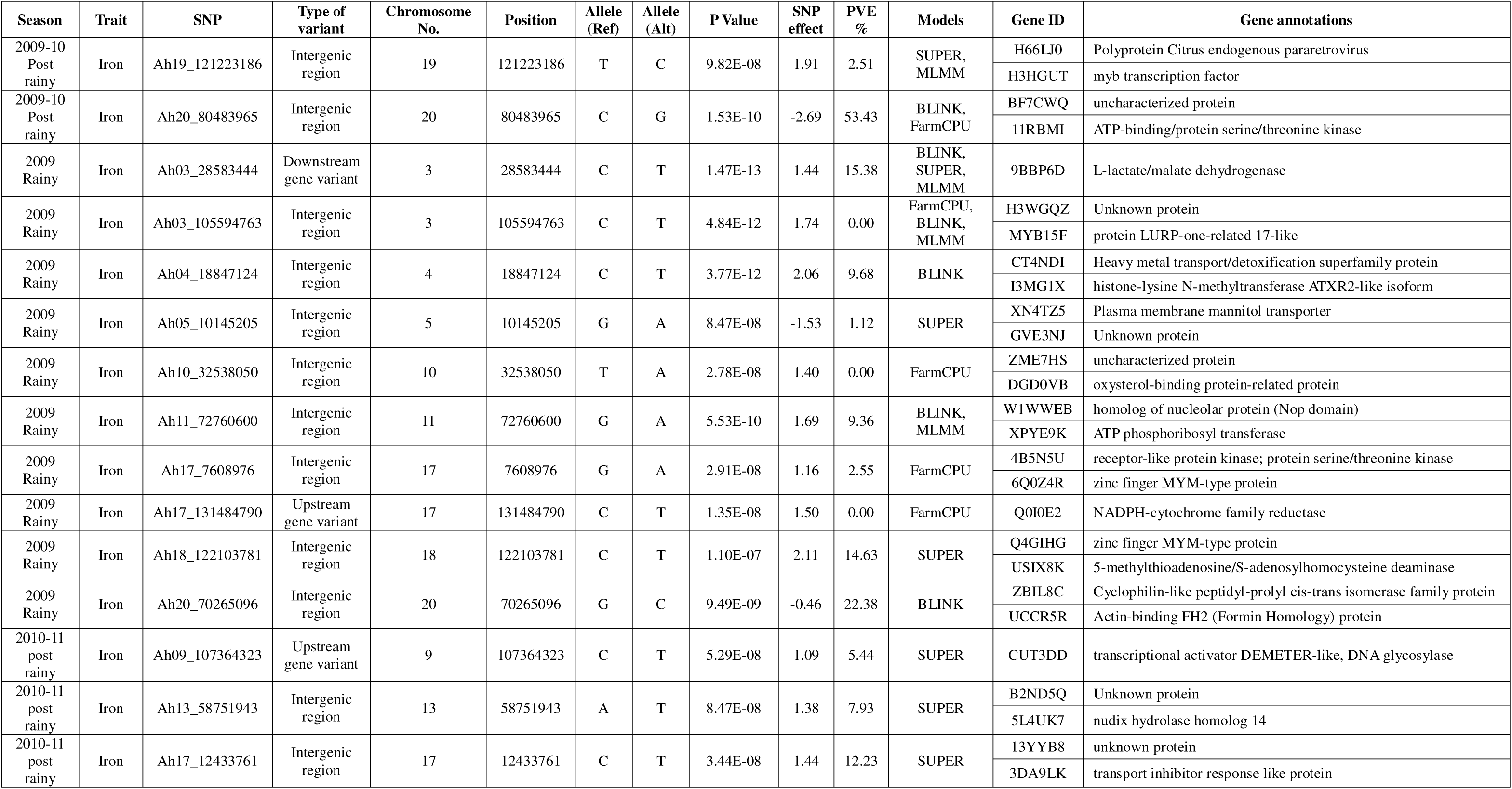
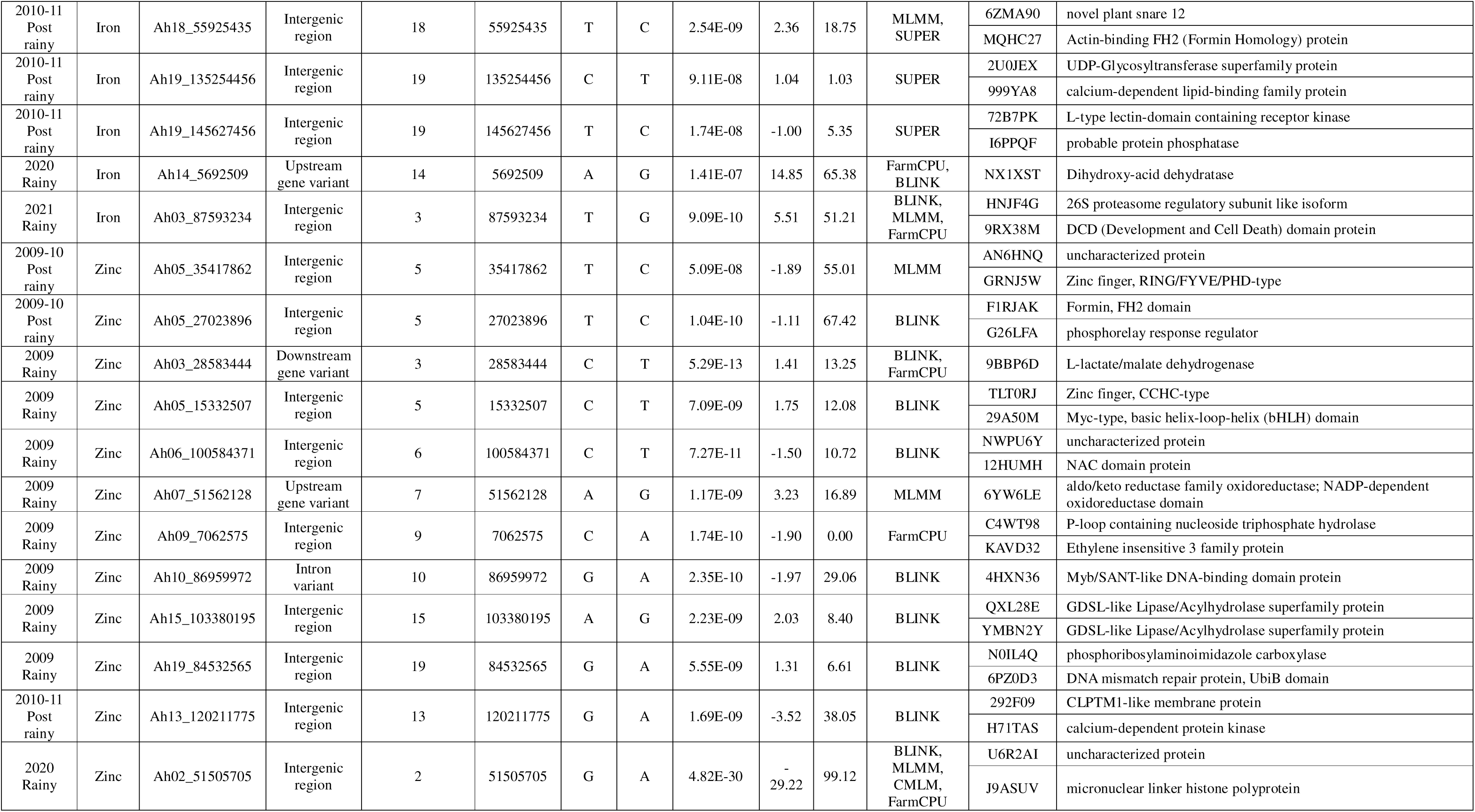
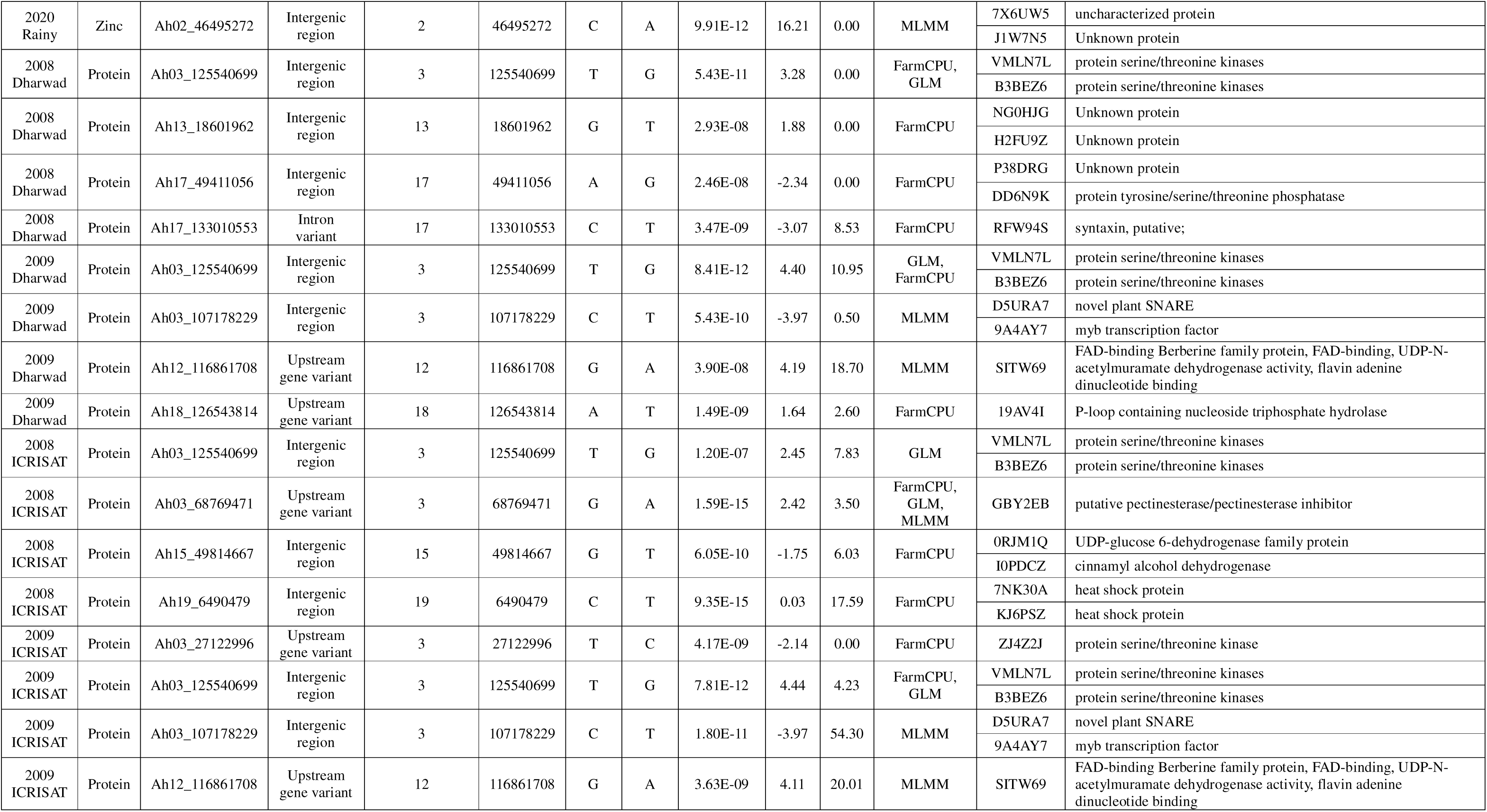
List of candidate genes in significant MTAs for Fe, Zn and PC using different models for season-wise data.

#### 3.2.1. Genomic associations identified for Fe content

A total of 4 significant MTAs were identified for Fe on chromosomes Ah02 (Ah02_5010027), Ah13 (Ah13_17399102), Ah19 (Ah19_27281401) and Ah20 (Ah20_80483965) (**Figure 2A**). The *p-values* for these significant SNPs were identified with the values ranging from 1.20 × 10^-8^ (Ah20_80483965) to 5.14 × 10^-8^ (Ah02_5010027) which explained 14.04 to 26.57% of phenotypic variation. The SNPs Ah13_17399102 and Ah20_80483965 were uniquely identified by BLINK model. Among the SNPs associated with Fe, the SNP Ah02_5010027 identified on chromosome Ah02 was detected by the models-BLINK, MLMM and SUPER. Ah02_5010027 was observed with the highest phenotypic variance of 26.57% with a ‘*p*’ value of 5.14 × 10^-8^ whereas Ah13_17399102 was found to have the lowest phenotypic variance of 11.27% with a ‘*p*’ value of 3.96 × 10^-8^. The SNPs, Ah19_27281401 and Ah20_80483965, located on chromosomes Ah19 and Ah20 were identified with phenotypic variance values of 18.18% and 14.04%, and ‘*p*’ values of 1.80 × 10^-9^ and 1.20 × 10^-8^, respectively.

**Figure 2.**
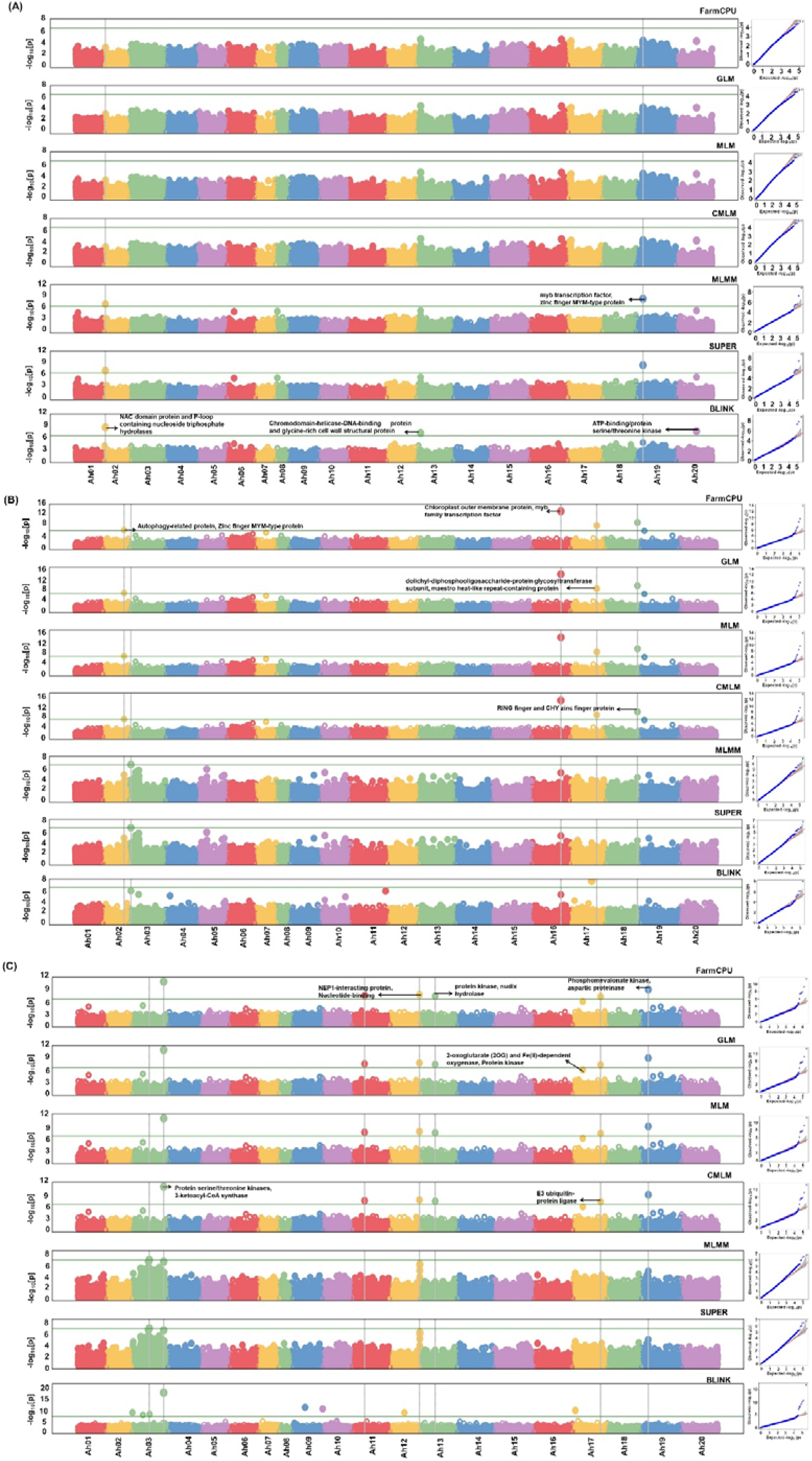
Manhattan plots and optimal Q-Q plots for (A) Fe (B) Zn and (C) PC of pooled season data. The horizontal dashed lines represent the significant threshold (*p* = 1 × 10^−6,^ Bonferroni correction) and the vertical dashed lines represent the candidate genes at the significant SNPs associated with (A) Fe content on chromosomes-Ah02, Ah13, Ah19 and Ah20. (B) Zn content on chromosomes-Ah02, Ah16, Ah17 and Ah18. (C) protein content on chromosomes-Ah03, Ah11, Ah12, Ah13, Ah17 and Ah19.

For seasonal analysis, 20 significant MTAs were identified for Fe on different chromosomes Ah03, Ah04, Ah05, Ah09, Ah10, Ah11, Ah13, Ah17, Ah18, Ah19 and Ah20. The *p-values* for these significant SNPs were identified with the values ranging from 1.10 × 10^-7^ (Ah18_122103781) to 1.47 × 10^-13^ (Ah03_28583444) which explained 14.63 to 15.37% of phenotypic variation. Three MTAs on chromosome Ah03 (Ah03_28583444, Ah03_105594763 and Ah03_87593234) were identified by at least three models during S2 and S5 (**Supplementary Figures S1, S2**). The highest phenotypic variance (65.38%) was observed in Ah14_5692509 with a ‘*p*’ value of 1.41 × 10^-7^ followed by Ah03_87593234 (51.20%) with a ‘*p*’ value of 9.09 × 10^-10^.

#### 3.2.2. Genomic associations identified for Zn content

Similarly, 4 significant MTAs were identified for Zn on chromosomes Ah02 (Ah02_80149889), Ah16 (Ah16_114034439), Ah17 (Ah17_102733285), and Ah18 (Ah18_127902116) (**Figure 2B**). All the SNPs associated with the Zn were detected by the models-FarmCPU, GLM, MLM and CMLM. The *p*-values for these identified SNPs were ranging from 3.28 × 10^-9^ (Ah17_102733285) to 1.19 × 10^-14^ (Ah16_114034439). Furthermore, Ah16_6885633 was observed to have highest phenotypic variance of 20.49% and Ah17_102733285 was observed to have lowest phenotypic variance of 0.71%. The SNPs, Ah02_80149889 and Ah18_127902116, located on chromosomes Ah02 and Ah18 were identified with phenotypic variance values of 14.83% and 12.48%, and ‘*p*’ values of 1.38 × 10^-7^ and 2.40 × 10^-10^, respectively.

Seasonal analysis identified 13 significant MTAs on chromosomes Ah02, Ah03, Ah05, Ah06, Ah07, Ah09, Ah10, Ah13, Ah15 and Ah19. The *p-values* for these significant SNPs were identified with the values ranging from 5.09 × 10^-8^ (Ah05_35417862) to 4.82 × 10^-30^ (Ah02_51505705) which explained 55.01% to 99.12% of phenotypic variation. The MTA Ah02_51505705 on chromosome Ah02 was detected by the four models (BLINK, CMLM, FarmCPU, MLMM) during S4 (**Supplementary Figure S3, S4**). The highest phenotypic variance (99.12%) was observed in Ah02_51505705 with a ‘*p*’ value of 4.82 × 10^-30^ followed by Ah05_27023896 (67.42%) with a ‘*p*’ value of 1.04 × 10^-10^.

#### 3.2.3. Genomic associations identified for PC

Total, 7 significant MTAs were identified for PC on six different chromosomes like Ah03 (Ah03_2532408), Ah11 (Ah11_48679516), Ah12 (Ah12_56160933), Ah13 (Ah13_58948078), Ah17 (Ah17_10878383 and Ah17_110193095), Ah19 (Ah19_28004976) (**Figure 2C**). All the SNPs associated with PC were detected by FarmCPU, GLM, MLM and CMLM models. The *p*-values for these SNPs encoding PC were ranging from 1.99 × 10^-8^ (Ah11_48679516) to 5.72 × 10^-10^ (Ah19_28004976), which explained 1.84% to 16.05% of phenotypic variation. Interestingly, two candidate genes coding for the PC were found on the chromosome Ah17 at two different positions. The highest phenotypic variance of 16.05% was observed for Ah03_125540699 with a ‘*p*’ value of 4.70 × 10^-12^ and the lowest phenotypic variance of 1.84% was observed for Ah13_58948078 with a ‘*p*’ value of 2.67 × 10^-8^.

A total of 11 significant MTAs were resulted in seasonal analysis on chromosomes Ah03, Ah12, Ah13, Ah15, Ah17, Ah18 and Ah19. The *p-values* for these significant SNPs were identified with the values ranging from 1.20 × 10^-7^ (Ah03_125540699) to 9.35 × 10^-15^ (Ah19_6490479) which explained 7.82% to 17.59% of phenotypic variation. The MTA Ah03_68769471 on chromosome Ah03 was detected by the three models (FarmCPU, GLM, MLMM) during 2009 rainy season in ICRISAT (**Supplementary Figures S5, S6**).

### 3.3. Identification of candidate genes and their expression analysis

A total of 28 candidate genes were identified for pooled season data, including 8 for Fe, 8 for Zn and 12 for PC and seasonal data analysis identified 62 genes, including 30 for Fe, 18 for Zn and 14 for PC, associated with the MTAs. These candidate genes were involved in different molecular, cellular, and biological functions related to the traits. These candidate genes encode various important proteins like *MYB transcription factor*, *Zn finger MYM type protein*, *RING finger MYM type protein*, *ATP binding/ threonine kinase/ protein serine/ protein kinase*, *NAC domain protein*, and *glycine rich cell wall structural protein*. Several other genes like *Heavy metal transport/detoxification superfamily protein*, *ATP phosphoribosyl transferase*, *NADPH-cytochrome family reductase*, *nudix hydrolase*, *Probable protein phosphatase*, *Calcium-dependent protein kinase*, *Heat shock protein*, *E3 ubiquitin-protein ligase*, *2-oxoglutarate* (*2OG*) and *Fe* (*II*)*-dependent oxygenase superfamily protein*, *phosphomevalonate kinase*, *aspartic proteinase*, *3-ketoacyl-CoA synthase*, *dolichyl-diphosphooligosaccharide-protein*, *jasmonic acid carboxyl methyltransferase* were identified as candidate genes related to the trait functions and biological processes. The domains for the proteins encoded by these genes were either directly or indirectly related to the Fe, Zn and PC. The SNPs, Ah02_5010027 associated with *NAC domain protein*, Ah02_80149889, Ah18_127902116, Ah13_17399102, Ah17_7608976, Ah18_122103781, Ah05_35417862, Ah05_15332507 associated with *RING and Zn finger protein*, Ah16_114034439, Ah18_27281401, Ah19_121223186, Ah10_86959972 associated with *MYB transcription* were found linked to Fe and Zn homeostasis pathway.

Among the total candidate genes identified, 17 genes showed expression during different developmental stages (**Figure 3**). Gene expression studies show that *NAC domain protein* (*Arahy.LV3APC*), *Heavy metal transport/detoxification superfamily protein* (*Arahy.CT4NDI*), *2-oxoglutarate* (*2 OG*) and *Fe* (*II*)-*dependent oxygenase* (*Arahy.KX8RC4*) and *Heat shock protein* (*Arahy.7NK30A*) were found to show higher expression in majority of the plant tissues studied, whereas *glycine rich cell wall structural protein* (*Arahy.LJ25B0*), *MYB transcription factor* (*Arahy.QI0PHV*) and *Phosphomevalonate kinase* (*Arahy.QVG73I*) showed higher expression in some specific plant tissues only.

**Figure 3.**
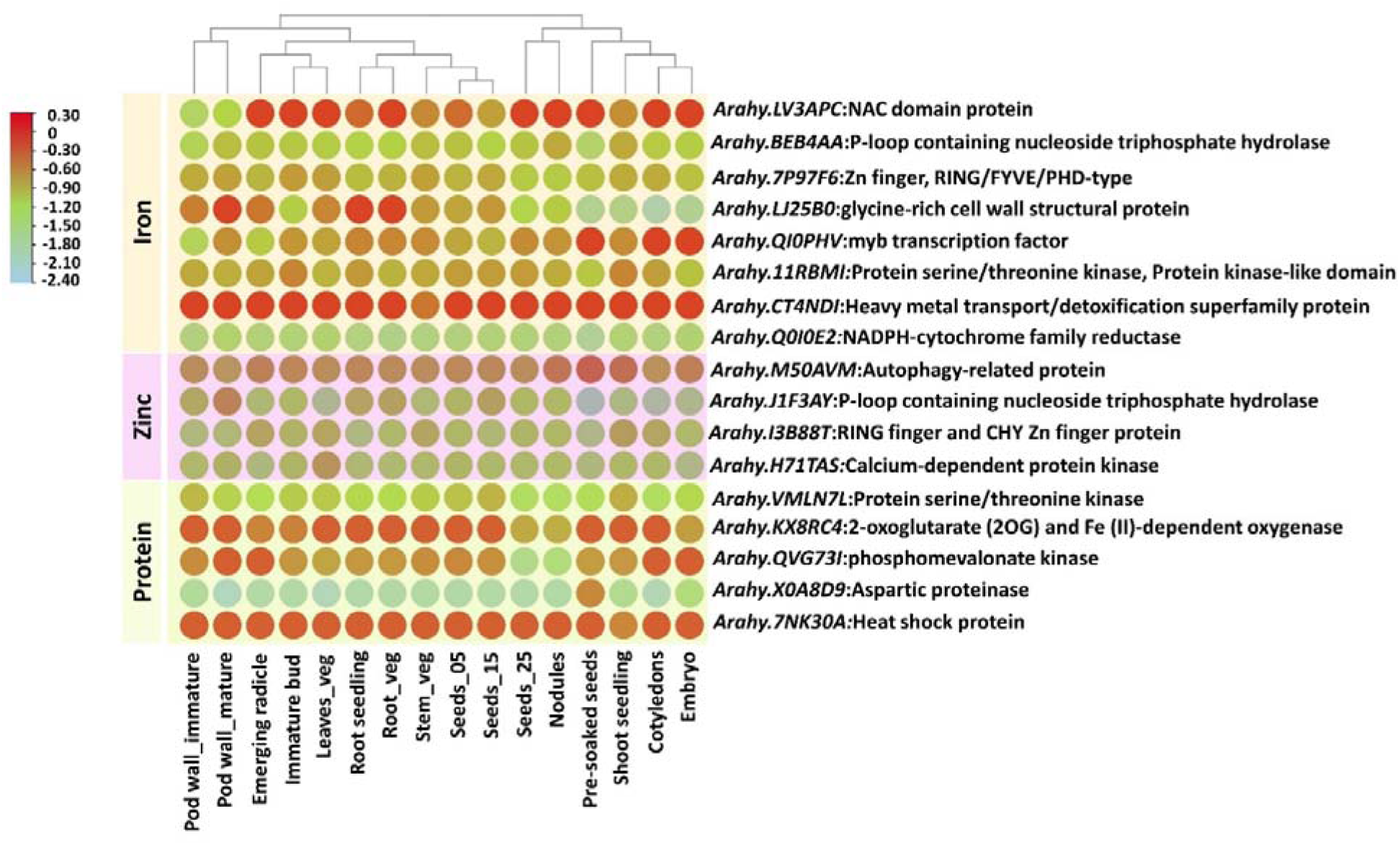
Tissue-specific expression study of candidate genes identified at significant MTA regions for Fe, Zn and PC using *Arachis hypogaea* gene expression atlas. The expression of 17 candidate genes in 16 different tissues (Pre-soaked seeds, emerging radicle, cotyledons, embryo, root seedling, leaves veg, root veg, shoot veg, shoot seedling, immature bud, seeds_05, seeds_15, seeds_25, pod wall immature, pod wall mature, nodules) are plotted in the heatmap.

### 3.4. Allele mining for significantly associated SNPs

In the current study, significant SNPs for all the three traits and the candidate genes associated with these genomic regions were obtained on different chromosomes. In order to use these genomic regions and candidate genes, allele call variation at the significant SNP position was carried out in a diverse mini-core collection of groundnut genotypes.

Based on the significant SNP positions associated with the three traits, allele mining was carried out to identify the favourable and un-favourable alleles in a diverse mini-core panel of groundnut genotypes (**Supplementary Figure S7**). The panel consisted of 16 genotypes for Fe, 22 genotypes for Zn and 18 genotypes for PC. Among the 16 genotypes for Fe, high Fe lines ranged from 28.7 to 52.5 ppm and low Fe lines ranged from 15.5 to 19.4 ppm. Among the 22 genotypes for Zn, high Zn lines showed a range of 40.1 to 49.2 ppm and low Zn lines showed a range of 19.8 to 27.9 ppm. Similarly, among the 18 lines for PC, high PC lines ranged between 26.8 to 28.6% and low PC lines ranged between 18.7 to 19.8%.

Among the four SNPs associated with Fe, three SNPs Ah02_5010027 (26.57% PVE with a ‘*p*’ value of 5.14 × 10^-8^), Ah13_17399102 (11.27% PVE with a ‘*p*’ value of 3.96 × 10^-8^) and Ah20_80483965 (14.04% PVE with a ‘*p*’ value of 1.20 × 10^-8^) showed clear difference between high and low Fe lines. Among the four SNPs identified for Zn, two SNPs Ah16_114034439 (20.49% PVE with a ‘*p*’ value of 1.19 × 10^-14^) and Ah18_127902116 (12.48% PVE with a ‘*p*’ value of 2.40 × 10^-10^) showed clear difference between the high and low Zn content lines. Similarly, among the seven SNPs identified for PC, two SNPs Ah03_125540699 (16.05% PVE with a ‘*p*’ value of 4.70 × 10^-12^) and Ah19_28004976 (4.17% PVE with a ‘*p*’ value of 5.72 × 10^-10^) showed clear difference between high and low PC lines. Interestingly, one genotype ICG 9249- with mean values of Fe-30.1 ppm, Zn-40.3 ppm and PC-27.3% showed favourable alleles for the high Fe, Zn and PC and four genotypes, ICG 2777 (Fe- 19.4 ppm, Zn- 27.9 ppm), ICG 6813 (Fe- 19 ppm, Zn- 26.3 ppm), ICG 5016 (Fe- 18.4 ppm, Zn- 25.7 ppm) and ICG 14834 (Fe- 15.5 ppm, Zn- 19.8 ppm) showed unfavourable alleles for low Fe, and Zn.

Likewise, in-silico validation of the SNPs obtained from the seasonal data analysis is done using diverse set of 40 genotypes for both Fe and Zn (**Supplementary Figure S8**). Among the 20 SNPs associated with Fe, 8 SNPs- Ah05_10145205, Ah18_122103781, Ah20_70265096, Ah18_55925435, Ah19_135254456, Ah19_145627456, Ah14_5692509, Ah03_87593234 showed clear difference between high and low Fe lines. Similarly, among the 13 SNPs associated with Zn, three SNPs- Ah05_35417862, Ah05_27023896 and Ah02_46495272 showed clear difference between high and low Zn lines. Seven high Fe and Zn lines- ICG 14118 (Fe- 42.9 ppm, Zn- 53.3 ppm), ICG 15309 (Fe- 37.6 ppm, Zn- 52.9 ppm), ICG 36 (Fe- 36.4 ppm, Zn- 56.6 ppm), ICG 4911 (Fe- 35.2 ppm, Zn- 53.9 ppm), ICG 7963 (Fe- 34.7 ppm, Zn- 47.7 ppm), ICG 10092 (Fe- 34 ppm, Zn- 48.4 ppm) and ICG 1274 (Fe- 31.1 ppm, Zn- 47.7 ppm) showed favourable alleles whereas the genotypes ICG 8760 (Fe- 20.9 ppm, Zn- 25.7 ppm), ICG 4412 (Fe- 20.6 ppm, Zn- 13.7 ppm), ICG 4156 (Fe- 20.3 ppm, Zn- 15.9 ppm), ICG 4389 (Fe- 19.7 ppm, Zn- 16.4 ppm), ICG 76 (Fe- 18.9 ppm, Zn-14.2 ppm), ICG 4527 (Fe- 17.7 ppm, Zn- 15.3 ppm), ICG 13723 (Fe- 10.9 ppm, Zn- 14.8 ppm) and ICG 5016 (Fe- 9.2 ppm, Zn- 24.4 ppm) showed unfavourable alleles for the SNPs associated with both Fe and Zn. These findings suggest that these markers could be effectively deployed in the allele-specific marker development for marker-assisted selection (MAS) in improving the nutritional quality in groundnut.

### 3.5. Development and validation of KASP markers

Based on the allele call variations, 9 SNPs (2 SNPs on Ah05, 2 SNPs on Ah18, 2 SNPs on Ah19, 1 SNP each on Ah03, Ah14 and Ah20) were used to develop KASP markers. These markers were successfully developed and validated on a panel of 50 genotypes, which comprised of lines from reference set and mini-core collection of ICRISAT (**Figure 4**). The validation panel comprised of wide range of Fe and Zn contents, with Fe ranging from 7.6 to 42.9 ppm and Zn ranging from 10.9 to 62.4 ppm. Of the nine markers validated (eight for Fe and one for Zn), three KASPs (snpAH00636-intergenic SNP between *Arahy.HNJF4G* and *Arahy.9RX38M* on chromosome Ah03, snpAH00641-intergenic SNP between *Arahy.2U0JEX* and *Arahy.999YA8* on chromosome Ah19, snpAH00644-intergenic SNP between *Arahy.AN6HNQ* and *Arahy.GRNJ5W*) showed clear polymorphism in the diverse germplasm. These 3 KASP markers can be used as diagnostic markers for their effective utilization in marker-assisted breeding for enhancing Fe and Zn contents in groundnut.

**Figure 4.**
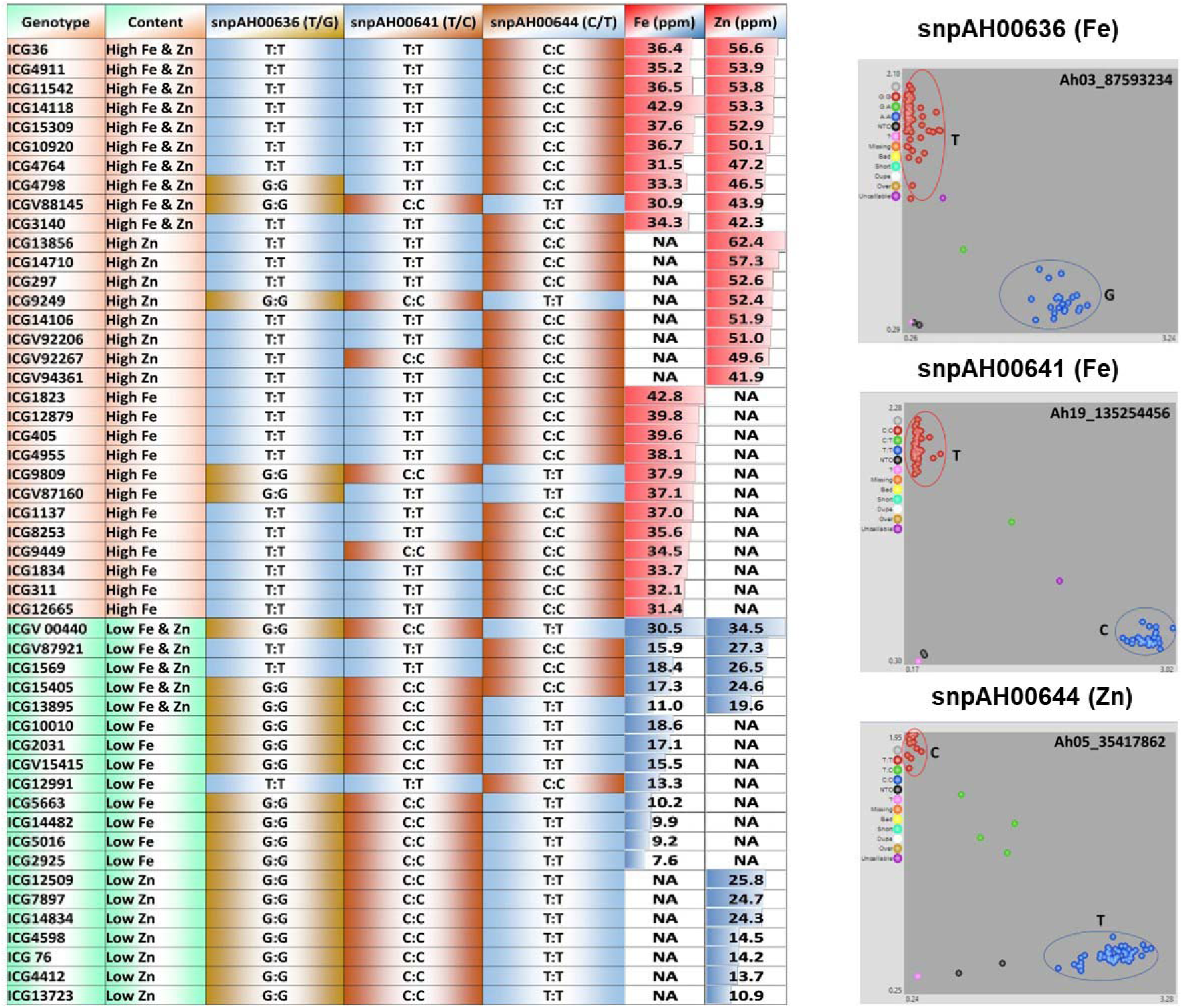
Validation of Kompetitive allele specific polymerase chain reaction (KASP) markers for Fe and Zn. KASP markers were validated in a panel consisting of 50 lines of both high and low Fe, Zn content lines with Fe ranging from 7.6 to 42.9 ppm and Zn ranging from 10.9 to 62.4 ppm. Three KASP markers (snpAH00636, snpAH00641 and snpAH00644) were successfully validated and show polymorphism between high and low Fe, Zn content genotypes. SNPs show clear homozygous clusters-snpAH00636 for genes *Arahy.HNJF4G* (*26S proteasome regulatory subunit*), *Arahy.9RX38M* (*Development and Cell Death domain protein*); snpAH00641 for genes *Arahy.2U0JEX* (*UDP-Glycosyltransferase superfamily protein*), *Arahy.999YA8* (*calcium-dependent lipid-binding family protein*); snpAH00644 for genes *Arahy.AN6HNQ* (*uncharacterized protein*), *Arahy.GRNJ5W* (*Zinc finger, RING/FYVE/PHD-type*).

### 3.6. Gene pathway for Fe and Zn homeostasis

The genes encoding the proteins and transcription factors associated with both Fe and Zn were found to suppress or inhibit the plant’s response to the deficiency conditions and also help in activation of a series of response mechanisms to enhance Fe and Zn uptake and homeostasis (**Figure 5**). These genes increase the uptake of Fe^+3^ and Zn^+2^ ions from the rhizosphere into the root tissues and also involve in the mobilization of these ions from root to the shoot and seed. The genes, *RING/ Zn finger domains* (*Arahy.7P97F6, Arahy.9R964H* and *Arahy.I3B88T*), which act as *E3 ubiquitin ligases* were identified to negatively regulate the Fe deficiency response. Similarly, *MYB 10* and *MYB 72* that belong to the family, *MYB transcription factor* (*Arahy.QI0PHV, Arahy.1I6ZSS*), induce the *NAS* genes under the Fe and Zn deficiencies that increase the levels of Fe and Zn in the shoot and the seed. Another transcription factor, *IDEF2* that belongs to the *NAC domain protein* family (*Arahy.LV3APC*), binds to cis-regulatory element, IDE2 present at the promoter site of Fe-deficiency response genes and controls the homeostasis of Fe by modulating the expression of these genes.

**Figure 5.**
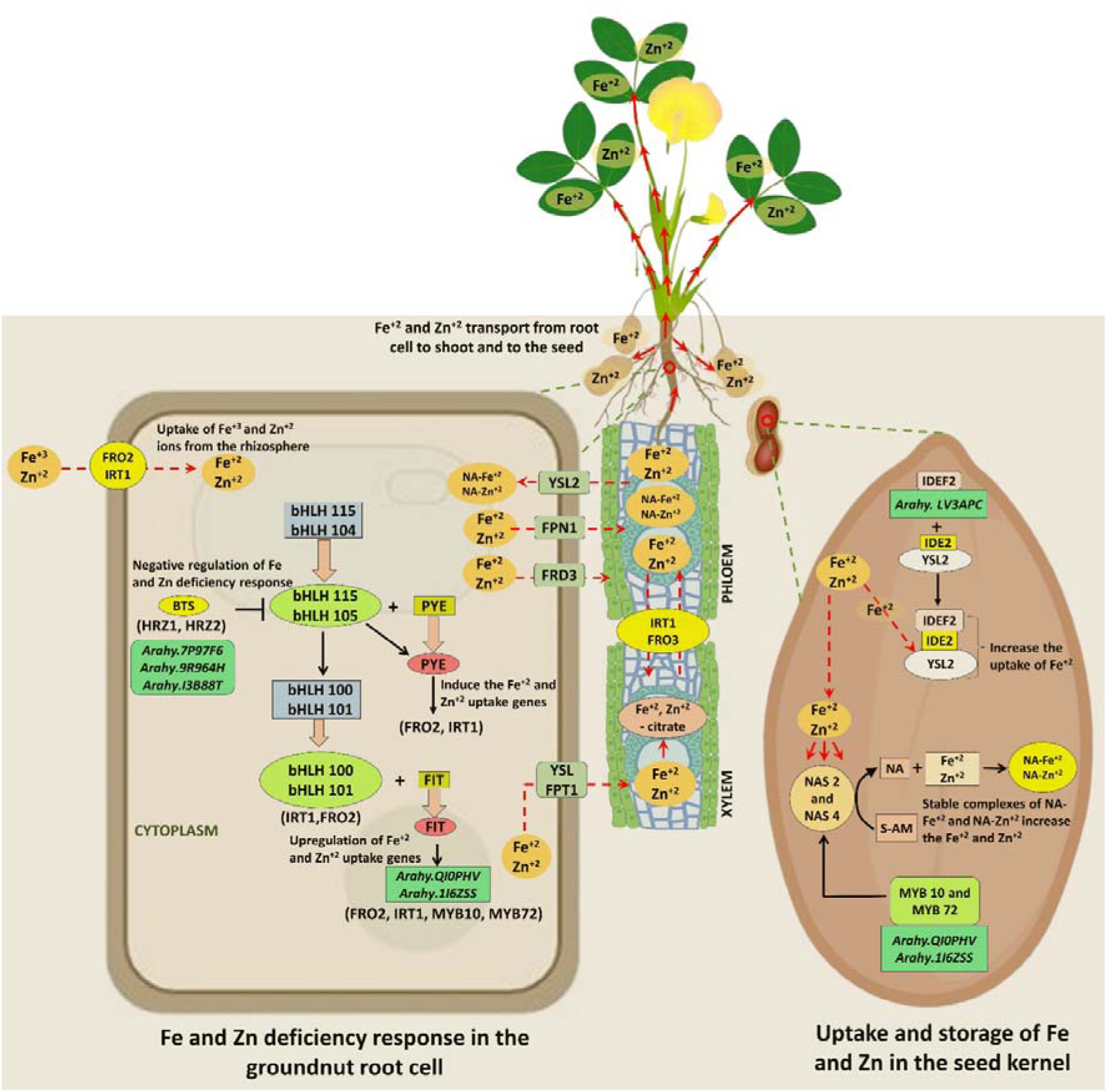
Molecular mechanism of Fe and Zn homeostasis in groundnut root cell and seed kernel. The figure depicts the Fe and Zn deficiency response in the groundnut root cell and the uptake, storage of both Fe and Zn in the groundnut seed kernel. bHLH – basic helix-loop helix; FIT– FER-like iron deficiency induced transcription factor; FRO- Ferric reduction oxidase; IRT- iron-regulated transporter; IRO- iron reduction oxidase; MYB- myeoloblastosis; PYE- POPEYE transcription factor; NAS- nicotianamine adenine synthase; HRZ- Haemerythrin motif-containing Really Interesting New Gene (RING)-and Zinc-finger protein; IDEF- iron deficiency responsive element binding factor; NA- nicotianamine adenine; YSL- yellow stripe like; VIT- vacuolar iron transporter; NRAMP- natural resistance associated macrophage protein; FRD- ferric reductase defective; FPN- ferriportin; IDE- iron deficiency responsive element; SAM- S-adenosyl L-methionine.

## 4. Discussion

A significant global health issue, malnutrition or hidden hunger, arises from the inadequate intake of nutrients in the diet. Biofortification offers a sustainable solution by improving crops with essential nutrients, thus linking agriculture directly to nutrition and health (30). Many crop varieties have been enriched with essential nutrients to combat malnutrition (30) including groundnut that offer essential nutrients and amino acids required for good health. Groundnut provides vital nutrients and amino acids crucial for optimum health, and further enhancement of its protein and micronutrient content could help address malnutrition. The detection and characterization of genomic regions and candidate genes related to these traits will advance biofortification efforts in groundnut.

In this study, the groundnut mini-core accessions were evaluated across five different seasons for the estimation of Fe and Zn, and across two locations for two seasons to estimate the PC. The phenotypic variability observed for these three traits was substantial, with similar variations reported for Fe, Zn and PC in groundnut (31), and other crops, such as chickpea (32) and wheat (33).

Multiple trait pyramiding in breeding can be achieved for studying correlated traits such as Fe and Zn (4). The significant positive correlation was observed between Fe, Zn and PC across locations and the seasons. Similar results were earlier reported in groundnut (*r* = 0.31 (4), *r* = 0.535 (31), *r* = 0.714 (15)) and other legume crops, including soybean (*r* = 0.29 (34)), mung bean (*r* = 0.469 (35)), and common bean (*r* = 0.416 (36)). These findings indicate that several common pathways have been associated with the homeostasis of Fe and Zn. In the PCA, Fe and Zn contributed positively towards the PC1 whereas, PC contributed positively towards both PC1 and PC2. Similarly, PC1 was positively influenced by Fe and Zn in groundnut and chickpea (37) and the contribution of both PC1 and PC2 was positively influenced by PC in chickpea (38).

GWAS analysis identified 15 MTAs for pooled data (4 for Fe, 4 for Zn, 7 for PC) and 44 MTAs for seasonal data (20 for Fe, 13 for Zn, 11 for PC). One stable MTA (Ah03_28583444) was detected on chromosome Ah03 for both Fe and Zn in S2. Similarly a previous study discovered a co-localized QTL linked with Fe and Zn on Ah03 (4). The genetic marker Ah03_28583444 closely associated with the MTA on chromosome Ah03 shows potential for further validation across wide range of populations and could be utilized for early-generation selection in breeding programmes (39). MTAs for both Fe and Zn were identified on chromosomes Ah02, Ah03, Ah05, Ah09, Ah10, Ah13, Ah17, Ah18, Ah19. In previous studies, significant QTLs for Fe and Zn were discovered on chromosomes Ah01, Ah03, Ah05, Ah07, Ah08, Ah11 and Ah13 (4). MTAs for PC were located on chromosomes Ah03, Ah11, Ah12, Ah13, Ah15, Ah17, Ah18, Ah19. Previous studies reported significant QTLs for PC on chromosomes Ah01, Ah03, Ah04, Ah05, Ah11, Ah12, Ah16, Ah18 (40). The highest number of MTAs for the three traits were identified on chromosomes Ah03 (8 MTAs), Ah17 (5 MTAs), Ah05 (4 MTAs) and Ah19 (4 MTAs). These genomic regions can be considered as hotspot regions which could help in the simultaneous improvement of the traits (41). One stable MTA (Ah03_125540699) associated with PC was identified on the Ah03 in four different seasons (2008R, 2009R of ICRISAT and 2008R, 2009R of Dharwad). Likewise, two stable MTAs (Ah03_107178229, Ah03_116861708) on chromosome Ah03 associated with PC were identified in two different seasons (2009R - Dharwad, 2009R - ICRISAT). Previous studies indicate that the environmental factors do not affect the expression of significant MTAs (42).

GWAS analysis in this study identified significant SNPs associated with the key candidate genes that regulate the Fe and Zn homeostasis pathways. These genes include *NAC domain protein* (*Arahy.LV3APC*), *RING and Zn finger protein* (*Arahy.7P97F6, Arahy.9R964H* and *Arahy.I3B88T*), and *MYB transcription factor* (*Arahy.QI0PHV, Arahy.1I6ZSS*). *Ferric Reduction Oxidase 2* (*FRO2*) present in the plasma membrane of root cell uptakes and reduces majority of the Fe found in the rhizosphere as low-soluble Fe^3+^ oxyhydrates (43) and *Iron-Regulated Transporter 1* (*IRT1*) transports Fe^2+^ across the membrane. *FER-like Iron deficiency-induced transcription factor* (*FIT*), which shows expression only in roots, upregulates the *FRO2, IRT1, MYB10* and *MYB72* genes involved in Fe and Zn uptake (44). *FIT* forms a heterodimer with the bHLH proteins like bHLH100 and bHLH101, in order to bind to the promoter sites of *IRT1* and *FRO2* (45). *POPEYE* (*PYE*), another *bHLH* protein, operates independently of *FIT* (46) and is regulated by dimers of bHLH104 and ILR3, to bind with the transcription factors, bHLH115 and ILR3 (bHLH105) (47).

According to recent studies, the Fe deficiency response may be negatively regulated a small family of *hemerythrin E3 ligases*, which includes *HRZ* in rice (48) and *BRUTUS* (*BTS*) in *Arabidopsis* (49). Interestingly, these hemerythrin domains of the *HRZ* proteins bind significant amounts of Zn along with Fe (48). *HRZ1* and *HRZ2* also have *Zn-finger domains* (*Arahy.7P97F6*, *Arahy.9R964H* and *Arahy.I3B88T*), *RING* (*really interesting new gene*) that function as *E3 ubiquitin ligases*.

The ferrous iron (Fe^+2^) is believed to be transported into the xylem via *Ferroportin1* (*FPT1*), and Zn^+2^ via YSL, once they enter the root, where these ions combine with citrate to create a complex, and then travels into the phloem where they combine with nicotianamine (*NA*) (50). *Nicotianamine synthase* (*NAS*) converts S-adenosyl methionine into NA, to create stable complexes with a variety of metals like Fe and Zn (51). Fe homeostasis is particularly dependent on NA-Fe^+2^ complexes since they are necessary for both remobilising Fe^+2^ and unloading the vasculature (52). Recent studies demonstrate that NA has a vital role in both Zn tolerance and hyperaccumulation in *Arabidopsis halleri*, underscoring its growing significance in mineral homeostasis and indicating possible applications of this molecule in bioremediation and biofortification (53). The ability of nicotianamine to bind to the divalent cations in the order Mn^+2^<Fe^+2^<Zn^+2^<Ni^+2^ with a high binding affinity is well-established (50). The limited amounts of NA will promote the development of NA-Zn^+2^ complexes over Mn^+2^ and Fe^+2^ in the case of limiting Fe conditions.

The genes *MYB transcription factor* (*Arahy.QI0PHV*, *Arahy.1I6ZSS*), and *NAC domain* (*Arahy.LV3APC*) were earlier reported to show their expression in groundnut seed kernel (54), (55). The *MYB family*, which consists of 126 members and is distinguished by the R2 and R3 MYB helix-turn-helix DNA-binding domains, includes *MYB10* and *MYB72* (*Arahy.QI0PHV, Arahy.1I6ZSS*) as close paralogs (56). Under Fe and Zn deficiency, *NAS2* and *NAS4* do not induce properly due to the absence of *MYB10* and *MYB72* (*Arahy.QI0PHV, Arahy.1I6ZSS*), which results in local decline in NA, reduced levels of NA bound Fe^+2^ and Zn^+2^, and eventually decreased levels of Fe and Zn in the seed and shoot. These findings suggest that *MYB10* and *MYB72* (*Arahy.QI0PHV*, *Arahy.1I6ZSS*) are essential for survival under Fe and Zn-limiting situations, for expression of both *NAS2* and *NAS4*, and to properly acquire Fe and Zn in plants (57). Plants develop a signalling network to regulate Fe uptake and transport in order to maintain Fe homeostasis. The transcription factor, *IDEF2* specifically binds to cis-acting element, IDE2 present at the promoter site of Fe deficiency-response genes in several plant species. *IDEF2*, a *NAC family* member (*Arahy.LV3APC)* controls Fe homeostasis by inducing the expression of a number of Fe deficiency-response genes (58).

This study also reported several other candidate genes that are either directly or indirectly associated with the functional regulation of Fe, Zn and PC in the plants. Previous studies have shown that *P-loop containing nucleoside triphosphate hydrolases* (*Arahy.BEB4AA*) are members of a unique class of molecules called metallochaperones, which play a key role in the nuclear and cytosolic compartmentalisation, storage, and export of Fe in the forms of ferritin and ferroportin, and are involved in Zn-ion binding (59). The gene*, ATP-binding/protein serine/threonine kinase* (*Arahy.11RBMI, Arahy.VMLN7L*, *Arahy.B3BEZ6*, and *Arahy.0IU8HI*), is involved in the homeostasis of Fe in *Arabidopsis* and the transcriptional activity regulation (60).

The *autophagy-related protein* (Arahy.*M50AVM*) is involved in the intracellular utilisation of Zn and inhibits the accumulation of reactive oxygen species (ROS) caused by Zn deficiency (61). *Calcium-dependent protein kinase* family helps in the stress responsive regulation of Zn^+2^ ions (62). The gene, *E3 ubiquitin-protein ligase* (*Arahy*.*PE3CF6*), identifies the target proteins and has crucial role in protein degradation and is a major component of ubiquitination cascade during plant response to stress (63). *Heat shock proteins* (HSPs) play an important role in protein folding, assembly, translocation, and degradation under stress and in several regular cellular activities (64).

Diagnostic markers enhance the development of improved varieties by enabling selection through marker-assisted early-generation screening, utilizing rapid generation advancement (65). The validated KASP markers for various traits like LLS, high oleic acid and leaf rust resistance, were widely utilized in groundnut breeding programs globally, contributing to the generation of thousands of advanced breeding lines, some of which have been made available for commercial cultivation (66). To identify and confirm the diagnostic markers associated with Fe and Zn contents, 9 SNPs were selected for the development of KASP markers, including SNPs on Ah03, Ah05, Ah14, Ah18, Ah19, Ah20. Primers were developed for the 9 SNPs and validated on a panel comprising diverse genotypes with Fe content ranging from 7.6 – 42.9 ppm and Zn content ranging from 10.9 – 56.6 ppm. Among the 9 KASP markers, 3 were successfully validated and showed clear differentiation between high and low Fe, Zn lines. Furthermore, these markers can be utilized to develop KASP assays and the genotypes carrying all the favourable alleles could be employed in marker-assisted breeding programmes for Fe and Zn biofortification.

The gene expression analysis showed that certain genes, such as *NAC transcription factor* (*Arahy.LV3APC), Heavy metal transport/detoxification superfamily protein* (*Arahy.CT4NDI*), *2-oxoglutarate and Fe (II)-dependent oxygenase* (*Arahy.KX8RC4*), *Heat shock protein* (*Arahy.7NK30A*) exhibit higher expression in multiple plant tissues, including seeds, roots, and nodules, whereas *MYB transcription factor* (*Arahy.QI0PHV*) exhibited higher expression in seeds, cotyledon and embryo, *Glycine-rich cell wall structural protein* (*Arahy.LJ25B0*) showed high expression in pod wall mature, root seedling and root veg. Earlier studies revealed the higher expression of *NAC genes* in root tissues, nodules, and developing seed (11), (55) and *MYB transcription factor* expressed highly in developing seed and shell (54), (11). This study demonstrates the crucial role of key candidate genes in Fe, Zn, and protein homeostasis. Identification and cloning of these key candidate genes will help in understanding the genetic and biological functioning of the traits. Additionally, identifying haplotypes for these genes using sequencing data from diverse germplasm may help in developing diagnostic markers for accelerated enhancement of the nutritional quality traits in groundnut.

## CONCLUSION

This study presents a comprehensive analysis of GWAS utilizing WGRS data and multi-season phenotypic data from a mini-core collection to identify significant MTAs and candidate genes related to Fe, Zn and PC in groundnut. A total of 15 MTAs and 28 candidate genes for pooled data, 44 MTAs and 62 candidate genes for seasonal data were identified for Fe, Zn and PC. Among the 9 KASP markers validated, three KASP markers (snpAH00636-intergenic SNP between *Arahy.HNJF4G* and *Arahy.9RX38M* on chromosome Ah03, snpAH00641-intergenic SNP between *Arahy.2U0JEX* and *Arahy.999YA8* on chromosome Ah19, snpAH00644-intergenic SNP between *Arahy.AN6HNQ* and *Arahy.GRNJ5W*) showed clear polymorphism between high and low content lines in the diverse germplasm. The key candidate genes including *MYB transcription factor* (*Arahy.QI0PHV, Arahy.1I6ZSS*), *Zn* finger *MYM type protein* and *RING finger MYM type protein* (*Arahy.7P97F6*, *Arahy.9R964H* and *Arahy.I3B88T*), *NAC domain protein* (*Arahy.LV3APC*) were found to be involved in the regulation of Fe and Zn homeostasis pathways. These candidate genes offer potential for haplotype analysis and the identification of the superior haplotypes, facilitating the generation of groundnut cultivars with enhanced nutritional value through haplotype-based breeding.

## Data availability statement

The sequencing data generated in this study is deposited in NCBI with bio-project ID (PRJNA1002116, PRJNA490835 and PRJNA490832).

## Institutional Review Board Statement

Not Applicable

## Informed Consent Statement

Not Applicable

## Author Contributions

MKP conceived the idea and supervised and finalized the manuscript. SP, RSB, AKP, KS phenotyped the mini-core collection. ND developed the hapmap file and mined gene sequences for marker development. UNS and SSG performed the analysis. UNS wrote the manuscript. UNS, DKM, SSG contributed in improvising figures. VS, KJ, MDS, PSR, RKV, MKP contributed to reviewing and improving the manuscript. All authors have read and agreed to the published version of the manuscript.

## Acknowledgments

The authors are thankful to GeneBank, ICRISAT for their support in providing seed material and assistance in phenotyping work. UNS is grateful to ICRISAT for providing the facilities to conduct the work.

