## Supplemental Figures and Tables for "Whole genome sequencing-based multi-locus association mapping for kernel iron, zinc and protein content in groundnut"

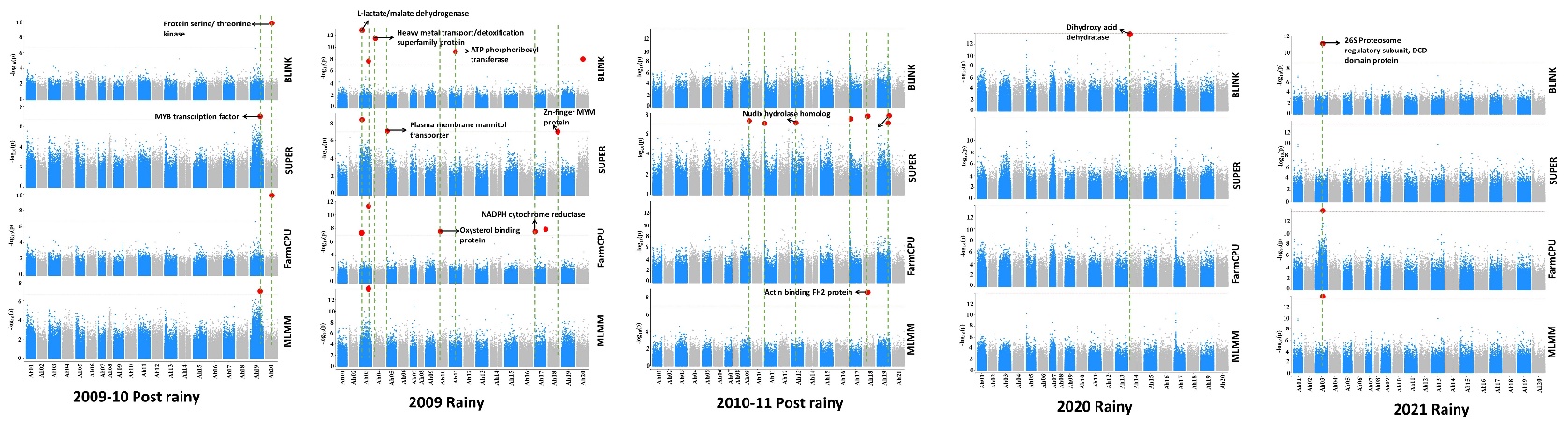
**Supplementary Figure S1** Manhattan plots showing significant MTAs for Fe content across five different seasons. The horizontal dashed lines represent the significant threshold kept at 6.7 level and the vertical dashed lines represent the candidate genes at the significant SNPs associated with Fe content.

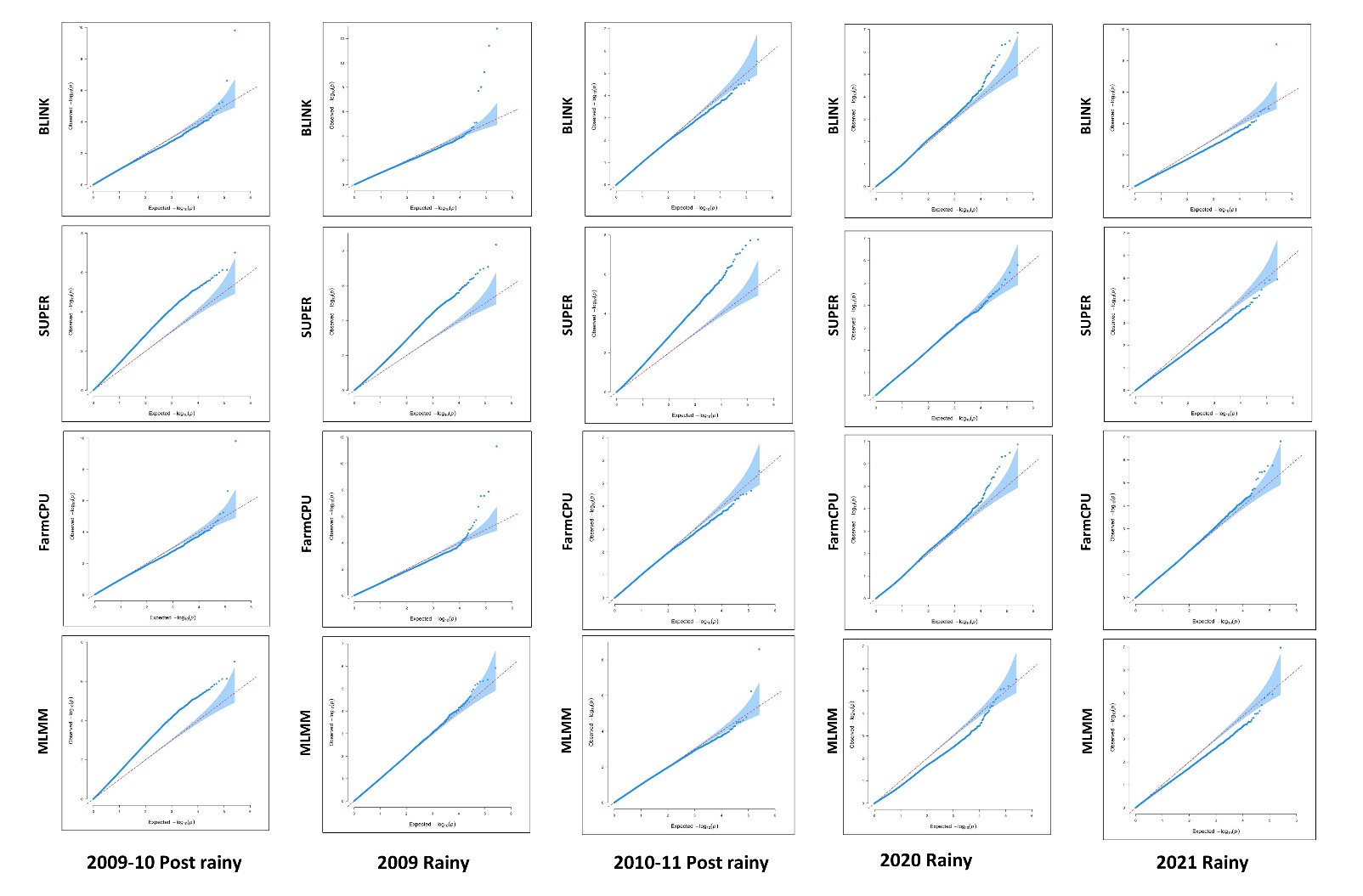
**Supplementary Figure S2** QQ plots for Fe content across five different seasons, to compare expected versus observed -log10(*p*) values.

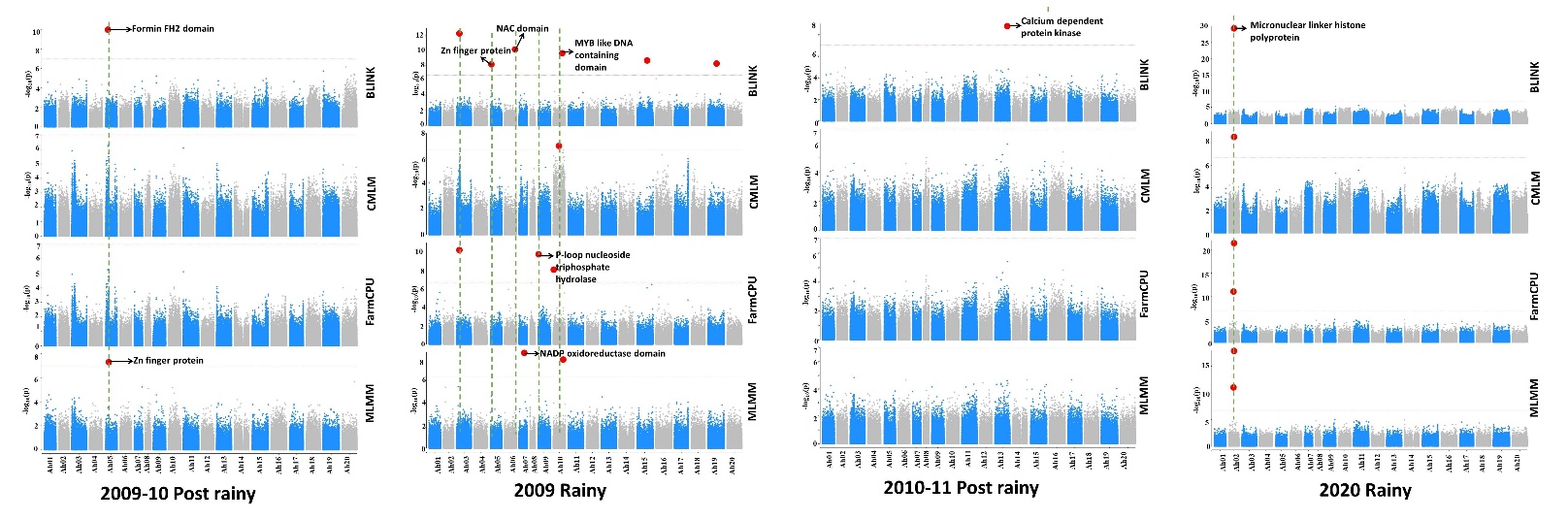
**Supplementary Figure S3** Manhattan plots showing significant MTAs for Zn content across four different seasons. The horizontal dashed lines represent the significant threshold kept at 6.7 level and the vertical dashed lines represent the candidate genes at the significant SNPs associated with Zn content.

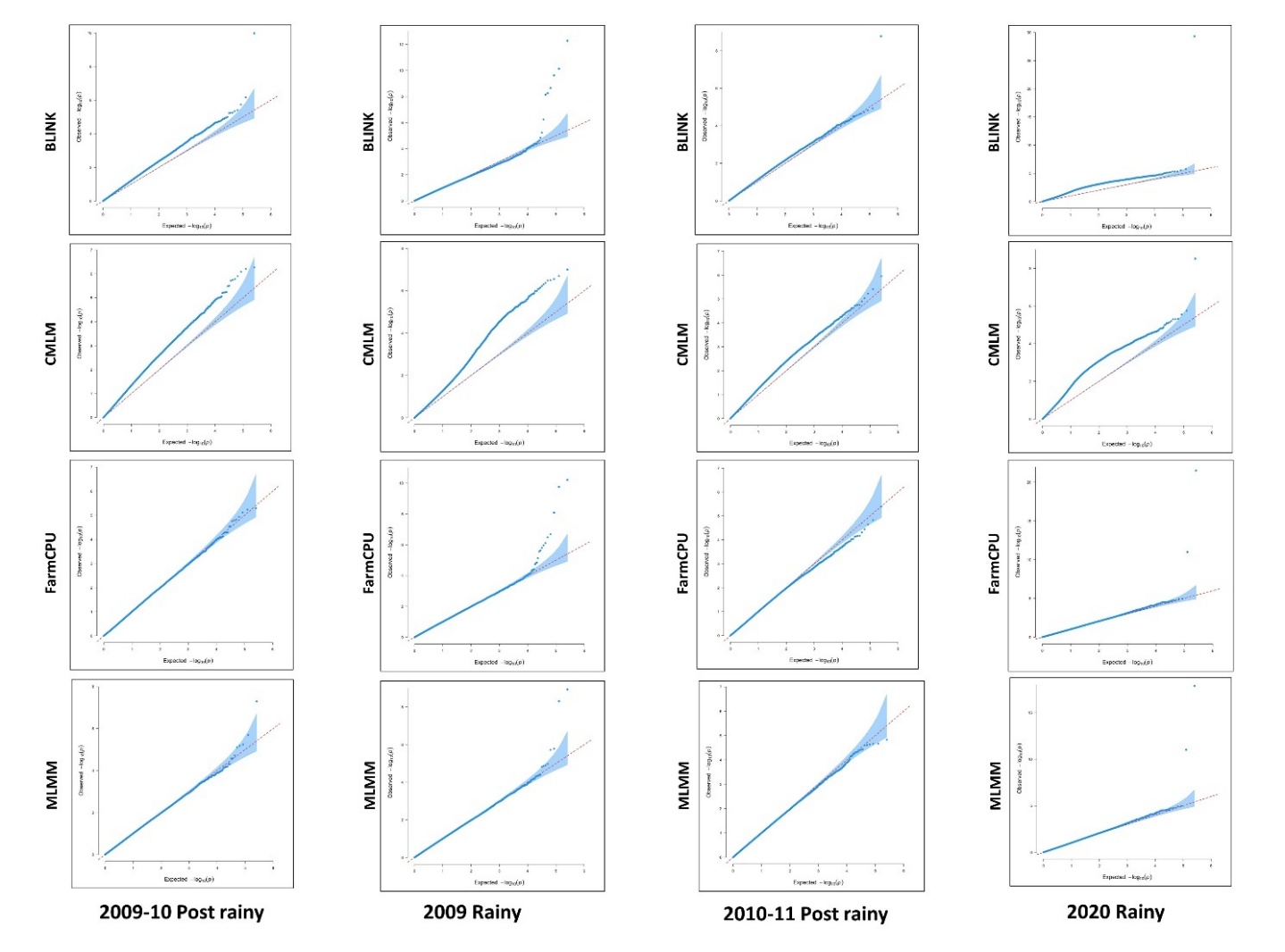

**Supplementary Figure S4** QQ plots for Zn content across four different seasons, to compare expected versus observed -log10(*p*) values.

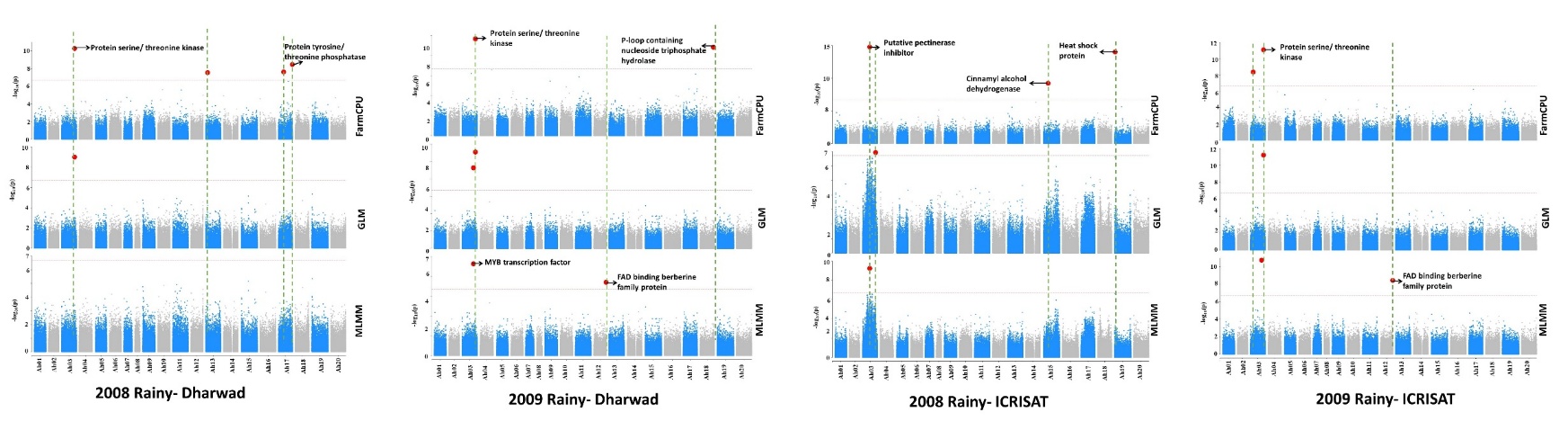
**Supplementary Figure S5** Manhattan plots showing significant MTAs for PC across four different seasons. The horizontal dashed lines represent the significant threshold kept at 6.7 level and the vertical dashed lines represent the candidate genes at the significant SNPs associated with PC.

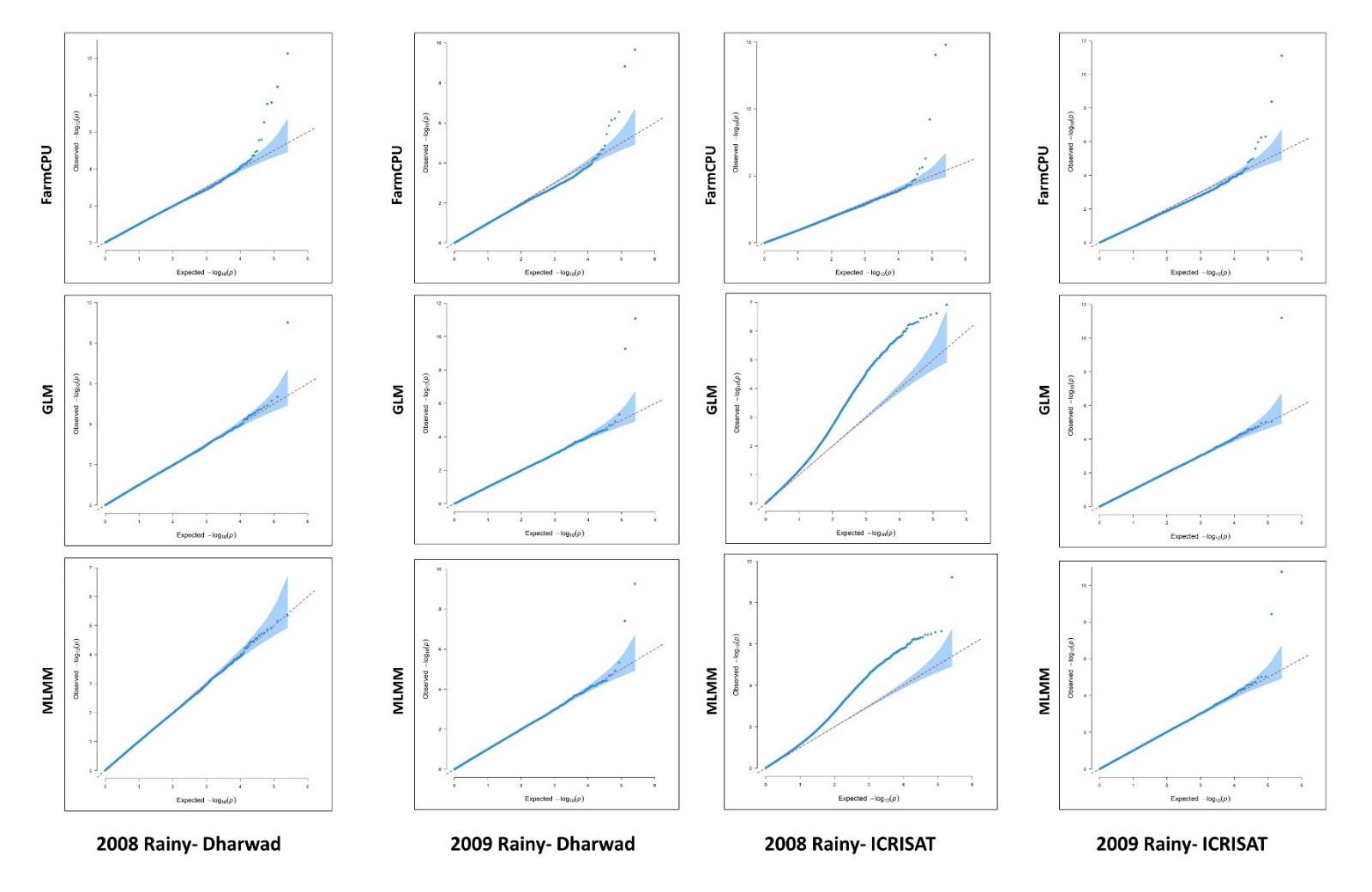

**Supplementary Figure S6** QQ plots for PC across four different seasons, to compare expected versus observed -log10(*p*) values.

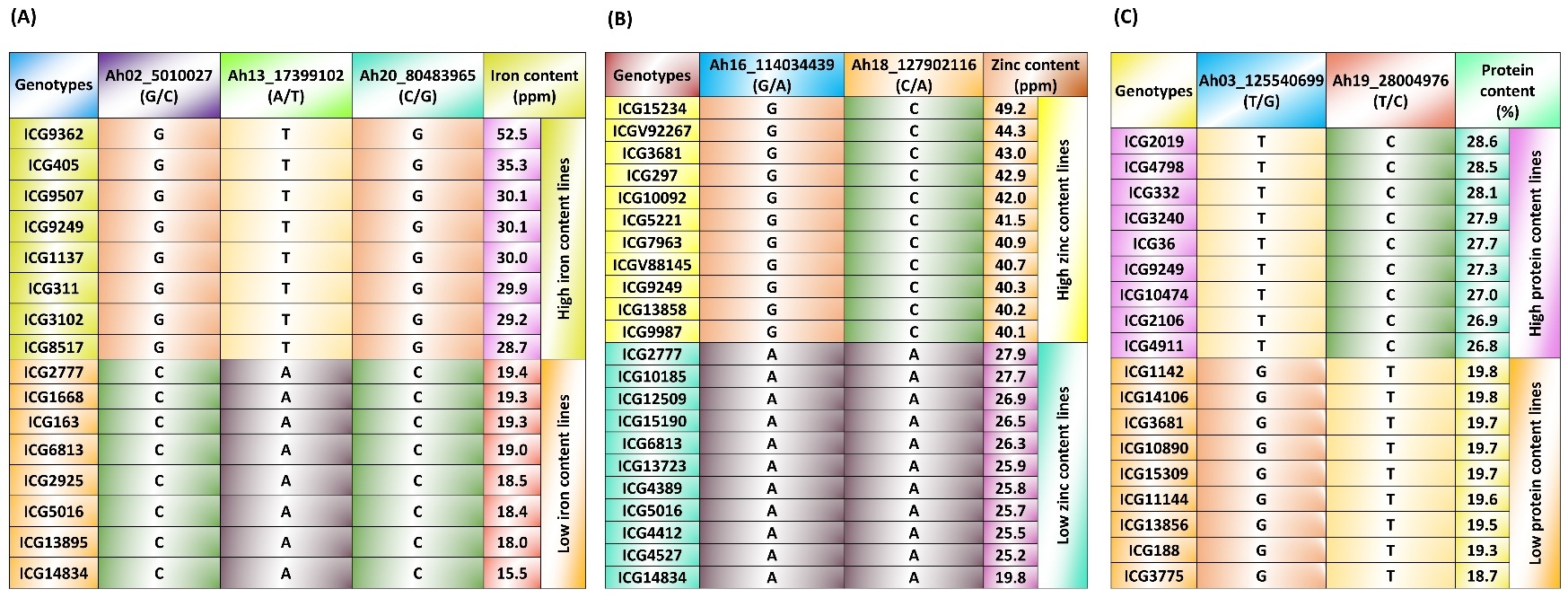

**Supplementary Figure S7** SNP call variations at the significant MTAs of pooled data analysis for three traits. Allelic variations among the (A) top 8 high Fe content lines and bottom 8 low Fe content lines, (B) top 11 high Zn content lines and bottom 11 low Zn content lines and (C) top 9 high PC lines and bottom 9 low PC lines -showing clear difference between the favourable and unfavourable allele calls at the significant SNP positions associated with the traits.

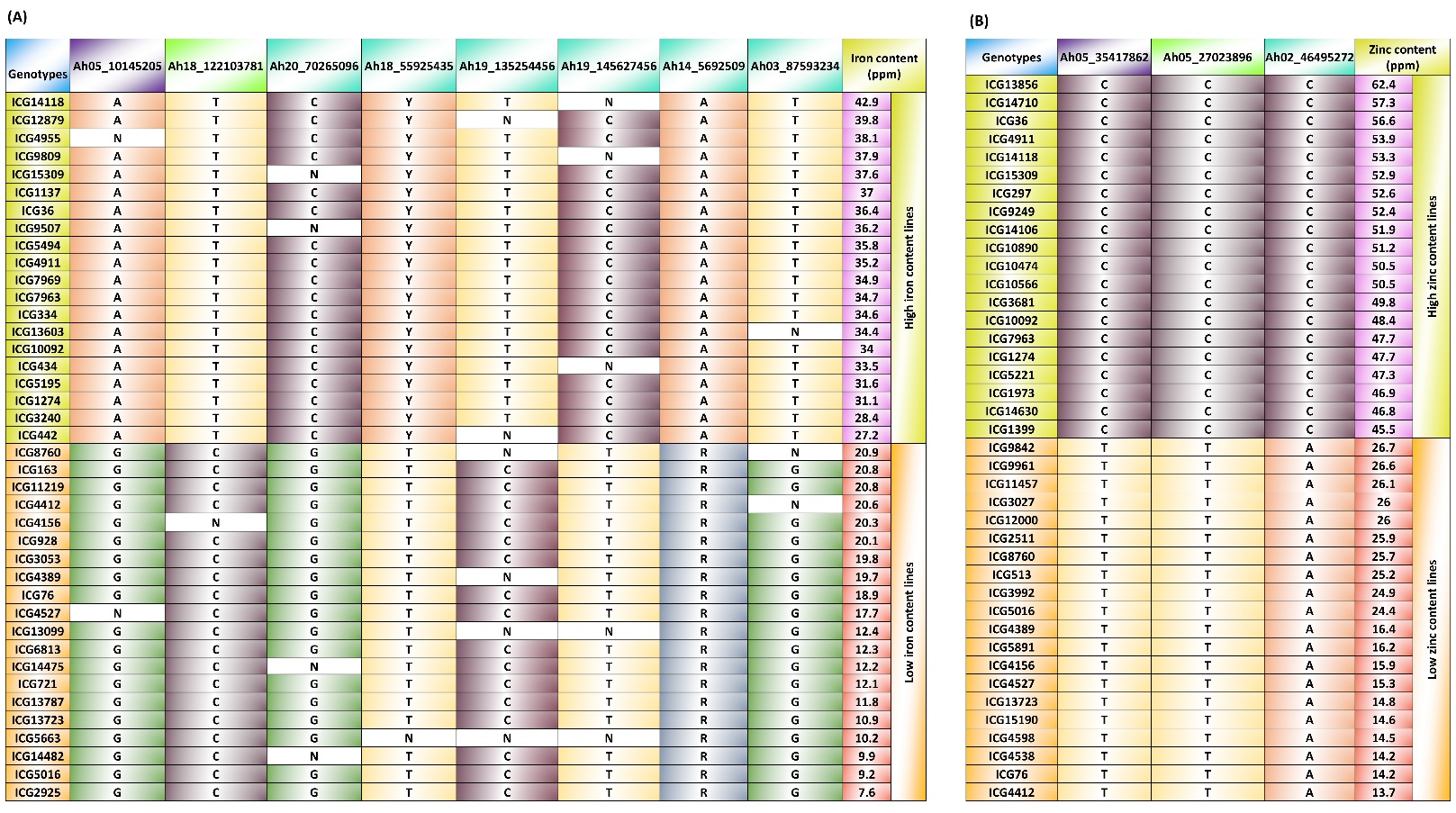

**Supplementary Figure S8** SNP call variations at the significant MTAs of seasonal data analysis for Fe and Zn. Allelic variations among the (A) top 20 high Fe content lines and bottom 20 low Fe content lines, (B) top 20 high Zn content lines and bottom 20 low Zn content lines -showing clear difference between the favourable and unfavourable allele calls at the significant SNP positions associated with the traits.

**Supplementary Table S1** Phenotypic data collected across the seasons and locations for Fe, Zn and PC

| Seasons  Traits | 2008 R | 2009 R | 2009-10 PR | 2010-11 PR | 2020 R | 2021 R |
| --- | --- | --- | --- | --- | --- | --- |
| Iron | - | L1 | L1 | L1 | L1 | L1 |
| Zinc | - | L1 | L1 | L1 | L1 | L1 |
| Protein | L1 & L2 | L1 & L2 | - | - | L1 | L1 |

L1: ICRISAT	L2: Dharwad	R: Rainy	PR: Post rainy

**Supplementary Table S2** Primer sequences for the three Kompetitive allele specific polymerase chain reaction (KASP) markers validated on diverse germplasm lines of groundnut

| **S. No.** | **Intertek ID** | **SNP name** | **Chromosome** | **Position** | **Reference allele** | **Alternate allele** | **Primer sequence** |  |
| --- | --- | --- | --- | --- | --- | --- | --- | --- |
| 1. | snpAH00636 | Ah03_87593234 | Ah03 | 87593234 | T | G | Forward | TGGGCCCCTTTGGGCCTCTTTTGTGGTCTTCGAGCTCTTGGAAAAGAGGTCGGTGGGGGACCAATCTTTCATAAGGTCGGTCCTTTTGTCTACTGCGAGTCTAACTCTGTAGCTCATGTCTGGGTATGAGCAGAGAGATTCACTCAAATTGAGGCACAAGGAAGGAAAGACAAGGAAGCTATCAAGGCTAGAATTGTATACTTGTGTAAGGAGATATGGAAGGAGAGAAACAAAGCAACTTTAAACAATGCAGAAGTAAATCTATATTCAGCTATCATAAGAGCGATAATCACATAAAAA |
|  |  |  |  |  |  |  | Reverse | AATTCAAAAATGCAAATGAGTTAGAAGACACAAGGTAATTAAAGGGATCAAGTCAAAGGAGGTACAAAAAGGTTACCTGGAGACCCCCTCCAGGGGTGTGGATAAAGGCGAACCTAGATGCAACATTCTTAAAAGACACAGTTGAAGGAGCAATTAGAATTATAATAAGGGAAAATTTAGGAAGGTTGGTCACAGGAACTGCAGAGAAGATTAACACGTCTTCATGCTTGACTACAGAAACATGTTGATAAATCCATATTTTACGATGATTTTTGAGTTGAAAAGTATAGAATTTATCAA |
| 2. | snpAH00641 | Ah19_135254456 | Ah19 | 135254456 | T | C | Forward | CCATAAAAAATAACCCCCATACAACTCTCTGTCACTAGGACCAAAGAAAGTATATTCAAGTATTGGGAAGAGAGACTTCAATTCATATGTGGGTAGAGCATTCAAATTCTCATTCCCGGGAGCTTGCATGCATAGAGTTGAAGGGACTTGTATGGTTACTCCTTAGTTATTGGAATGTGTAAGCATGTAGTGTGCATGGCTTCCTCCTTTGTTTGTGCATCTTTCTCTTCTTCCATGTGTGAGGTGGAAAGTTCTCTAATTTACTAGAGTATGAACTCCTTTGATTAATTTTCTCACATT |
|  |  |  |  |  |  |  | Reverse | TTCCGTAAGATCTTGCATTGTATCTCTTGGAAAAGAATTTGCTTGTTTCTCTATCATATGCTTGGTAATCAATTCCACGTGTTTTTCTATATTTTTTATGCTACCTCAAGTGGGTAGAGTAAACTCAGCAATATTTTGAGTAAACTCAGCAAAAGTATTCTCTAGGTTTTTCAATGCCAGCTCAACAGATGATAGTTCTTAATAGAAATAAAGAGATGATGTTTTTTTTATAAGAATAAAAAGGCGATGGTTCCTGATATGAACAGAGAGCTGATGGTTCTTGGTATAAATAAGAAAATG |
| 3. | snpAH00644 | Ah05_35417862 | Ah05 | 35417862 | C | T | Forward | TGTACGCTTTTTCTTCTTTTTACATACACTTCTACTTCTTTTTCTTCTTCTTTTCTTATTTTTCTCTTATCTTTTTTTTCATTATTGTCGCCGTCATCACTACTACCACCACCTCCTCCTCCCTTTCCTTCTTTTTTCATTTGAATTTCTTCTTCTCATTCATTTTCCTCCTCCTTCTCATCCATCTTCATCATCACTGTTATCGTTATTGTCATCATTGTCGTCTTCTTCTTATACGTATAATAACATTTCTGTGTTTTCTAGTATTTGTGTGCTAAATAACACCAGAACTACAATTAT |
|  |  |  |  |  |  |  | Reverse | ATGCATGAACTAATTACACCAAATCCAAGTCAATGAGACTAAATGTACTATATTTAAATTCAAAATGCATCGAAATTACTTAATGATGACGACGCACAAATAAATTTAGTTCAAAACAAGTACAAACCAAGTACACCTTATTTAAATTCAGAATTCAGCGAAATTAATTAATGATGCCGACACACAAGCAAACTCAGTTCAAAACAAGCACATAAACTAGATTCAATCATCCACAAAAATACATACAAGTAAATTTTAAATCCAGAAAAGTCATTAATATGACTTAAATGACACCAGAAA |
